# Latent Regulatory Programs Generate Synthetic T Cell States with Enhanced Therapeutic Potential

**DOI:** 10.64898/2026.01.07.698190

**Authors:** Brandon M. Pratt, Genevieve N. Mullins, Nolan Brown, William D. Green, Jennifer Modliszewski, Fucong Xie, Lara van Rooyen, Vasyl Zhabotynsky, Alexander N. Jambor, Gabrielle Cannon, Jarred M. Green, Andrew Kennedy, Coral del Mar Alicea Paunto, Huitong Shi, Emily Merritt, Takeshi Egawa, Ashwin Somasundaram, Wei Wang, Jessica E. Thaxton, Gianpietro Dotti, H. Kay Chung, J. Justin Milner

## Abstract

Transcription factors (TFs) govern cell fate through coordinated gene-regulatory networks, yet the full potential of these networks to generate non-native, therapeutically advantageous cell states *in vivo* remains largely unexplored. We hypothesized that systematic gain-of-function (GOF) overexpression of TFs in CD8⁺ T cells, central mediators of immune protection, could reveal latent, or “hidden,” regulatory programs capable of generating synthetic T cell states with therapeutic utility. To test this, we developed single-cell GOF sequencing (scGOF-seq), a multiplexed platform for unbiased, *in vivo* mapping of GOF effects on T cell fate in immunocompetent mouse models of infection and cancer. scGOF-seq uncovered unexpected regulators of T cell differentiation and accumulation, including SOX2, OCT4, and GATA2, which are normally silenced during T cell differentiation. Notably, outside its native regulatory context, supraphysiologic cMyc GOF reprogrammed CD8⁺ T cells into a synthetic stem-effector hybrid state, enabling >5,000-fold antigen-dependent expansion and antitumor activity, contrasting sharply with its native function in driving terminal differentiation. scGOF-seq further identified TF modules that cooperate with cMyc GOF to promote robust CD8⁺ T cell responses in solid tumors. Together, these findings establish GOF perturbation as a powerful strategy for revealing latent immune regulatory programs and engineering synthetic immune states with therapeutic potential.

**One-Sentence Summary:** *In vivo* single-cell gain-of-function screening reveals latent transcriptional programs that can reprogram T cells into highly functional synthetic states.

## Main Text

Cell states represent a spectrum of phenotypes largely shaped by coordinated transcription factor (TF) activity (*1, 2*). In the immune system, CD8⁺ T cells undergo extensive differentiation, giving rise to functionally distinct states across diverse contexts, including viral infection and cancer (*3–6*). These differentiation programs are governed by TF networks operating within tightly constrained regulatory architectures shaped by signaling-dependent induction, transient expression dynamics, and chromatin-encoded feedback mechanisms(*7–11*). Redirecting CD8⁺ T cell differentiation toward therapeutically favorable states is therefore a central goal of T cell-based immunotherapy, motivating increasingly sophisticated genetic engineering strategies. To date, much of our understanding of T cell fate control derives from loss-of-function (LOF) approaches, including conditional knockout models, CRISPR-based perturbations, and RNA interference(*11–14*). These studies have identified essential regulators of differentiation and enabled targeted enhancement of T cell function(*12, 14–16*). However, LOF strategies inherently define what a cell requires rather than the full range of achievable cell states. In addition, depleting a single TF often triggers regulatory compensation and plasticity, which can obscure latent regulatory potential (*17*).

In parallel, the clinical success of adoptive T cell therapies in hematologic malignancies (*18, 19*) has expanded the genetic design space available for therapeutic engineering. Unlike endogenous immune responses, adoptive cell therapy permits extensive *ex vivo* manipulation, enabling gain-of-function (GOF) perturbations and synthetic biology approaches that are not constrained by physiological regulatory limits. Recent GOF screens have begun to identify regulators that enhance T cell activity in cancer (*20–28*), demonstrating the promise of enforced gene activation as a complementary strategy to LOF. Nevertheless, how GOF perturbations reshape the full landscape of CD8⁺ T cell states *in vivo* and whether they can systematically uncover non-native, therapeutically superior differentiation programs remains poorly understood.

Mechanistically, GOF perturbations can sustain TF activity beyond endogenous transcriptional and epigenetic feedback, bypassing redundant regulatory circuits and exposing latent biology inaccessible to LOF genetics. Indeed, individual GOF studies have revealed previously unappreciated regulators of T cell differentiation and function, such as BATF3 and LTBR (*20, 24, 29*). However, unbiased, single-cell-resolved interrogation of GOF perturbations in antigen-specific T cells *in vivo* has remained limited, due in part to immunogenicity constraints of transduced donor cells in syngeneic hosts and technical challenges in deconvolving GOF effects at cellular resolution(*30–32*). We therefore reasoned that systematic, enforced GOF perturbation of TFs in antigen-specific CD8⁺ T cells would uncover latent regulatory programs, promote the emergence of synthetic cell states, and reveal new avenues for therapeutic T cell reprogramming.

Guided by this rationale, we developed a GOF screening framework to uncover the synthetic potential of T cells by identifying regulators that confer therapeutic attributes absent from natural immune responses, while avoiding dependence on local chromatin accessibility or large, immunogenic CRISPRa-based systems. Inspired by prior *in vitro* GOF screening(*20*), we engineered single-cell GOF sequencing (scGOF-seq), a platform for systematic profiling of GOF perturbations in antigen-specific CD8⁺ T cells across immune competent infection and cancer models. scGOF-seq enables rigorous pairwise comparisons and large-scale, multiplexed *in vivo* mapping of T cell differentiation with single-cell resolution. To support this approach, we developed a non-immunogenic, MSCV-based retroviral open reading frame (ORF) library that achieves robust transduction of mouse T cells, stable *in vivo* expression, and compatibility with established disease models. These design features preserved library representation *in vivo*, resolved each GOF perturbation at single-cell resolution, and enabled multiplexed evaluation of TF GOF perturbations within the same biological context.

To systematically interrogate latent biological potential via scGOF-seq, we constructed ORF libraries encoding two classes of TFs: (i) TFs that are differentially expressed during T cell differentiation, and (ii) putative “hidden drivers” of differentiation, TFs that are typically silenced or minimally expressed in T cells. Using scGOF-seq, we uncovered previously unappreciated GOF activities of multiple TFs, including hidden drivers such as SOX2, OCT4, and GATA2, in modulating CD8⁺ T cell differentiation *in vivo*.

In particular, we discovered an unexpected role for cMyc, classically associated with T cell activation and early effector commitment(*33*), in governing T cell exhaustion and stemness. Upon forced expression outside its native physiological patterns, cMyc GOF displayed a context-dependent ability to sustain stemness programs and counter exhaustion-associated transcriptional networks. Enforced cMyc GOF induced multimodal reprogramming of T cell metabolism, proliferation, tissue localization, protein translation, and differentiation, enabling T cells to acquire synthetic properties not observed in endogenous or control cells during settings of persistent antigen. Building on these insights, we applied specific GOF perturbations, including cMyc alone or in combination with cooperating factors, to engineer T cells with improved persistence and functional resilience in solid tumors. Together, these findings introduce a framework for interpreting GOF effects within established CD8⁺ T cell states and demonstrate how enforced supraphysiologic TF activity can be leveraged to generate therapeutically advantageous, synthetic cell states.

### scGOF-seq maps the latent regulatory landscape of CD8⁺ T cells *in vivo*

Despite advances in multiplexed GOF genetic screening technologies(*20–27*), systematic interrogation of GOF phenotypes *in vivo* remains limited, particularly in immunocompetent settings. As a result, the capacity of enforced gene activity to rewire CD8⁺ T cell fate beyond native regulatory constraints remains poorly defined. While some GOF effects are predictable, such as reinforcement of stem-like features by TFs endogenously enriched in stem-like T cells, including TCF1 or FOXO1(*34, 35*), GOF perturbations are not restricted to amplifying existing programs. By driving gene activity beyond physiological levels and outside endogenous regulatory feedback, GOF can expose latent or “*synthetic”* regulatory functions that cannot be inferred from LOF studies or expression patterns alone(*24*). Defining these effects requires unbiased approaches capable of resolving how enforced gene activity rewires cell fate *in vivo*.

TFs are especially well-suited for probing this latent regulatory space, as enforced expression can impose new gene-regulatory architectures and redirect cell identity. GOF perturbations may be especially informative for TFs that are lowly expressed, transiently induced, or epigenetically silenced during T cell differentiation, yet may retain regulatory capacity when forcibly activated. To identify such candidates, we integrated protein-coding expression data from the Human Protein Atlas, bulk RNA-seq data, ImmGen, and high-coverage single-cell RNA-seq datasets (∼80,000 reads per cell) from murine infection and cancer models(*36–38*). This analysis revealed that a substantial fraction of annotated TFs are expressed at trace or relatively low levels in CD8⁺ T cells **(Fig. 1A,B; fig. S1A,B)**. Many of these lowly expressed TFs are linked to non-immune lineage specification or cellular processes not associated with T cell biology, underscoring their potential to impose alternative gene-regulatory architectures when overexpressed. Notably, many retain accessible DNA-binding motifs across CD8⁺ T cell chromatin landscapes **(Fig. 1C; fig. S1C, S2A)**, raising the possibility that enforced expression could engage dormant regulatory circuitry. Based on these observations, we selected TFs for GOF screening from two categories: (i) differentially expressed TFs (including TFs with established roles in T cell differentiation), and (ii) putative “hidden driver” TFs (*39*) that are minimally expressed or silenced in T cells but likely retain latent regulatory capacity **(Fig. 1C)**.

**Fig. 1.**
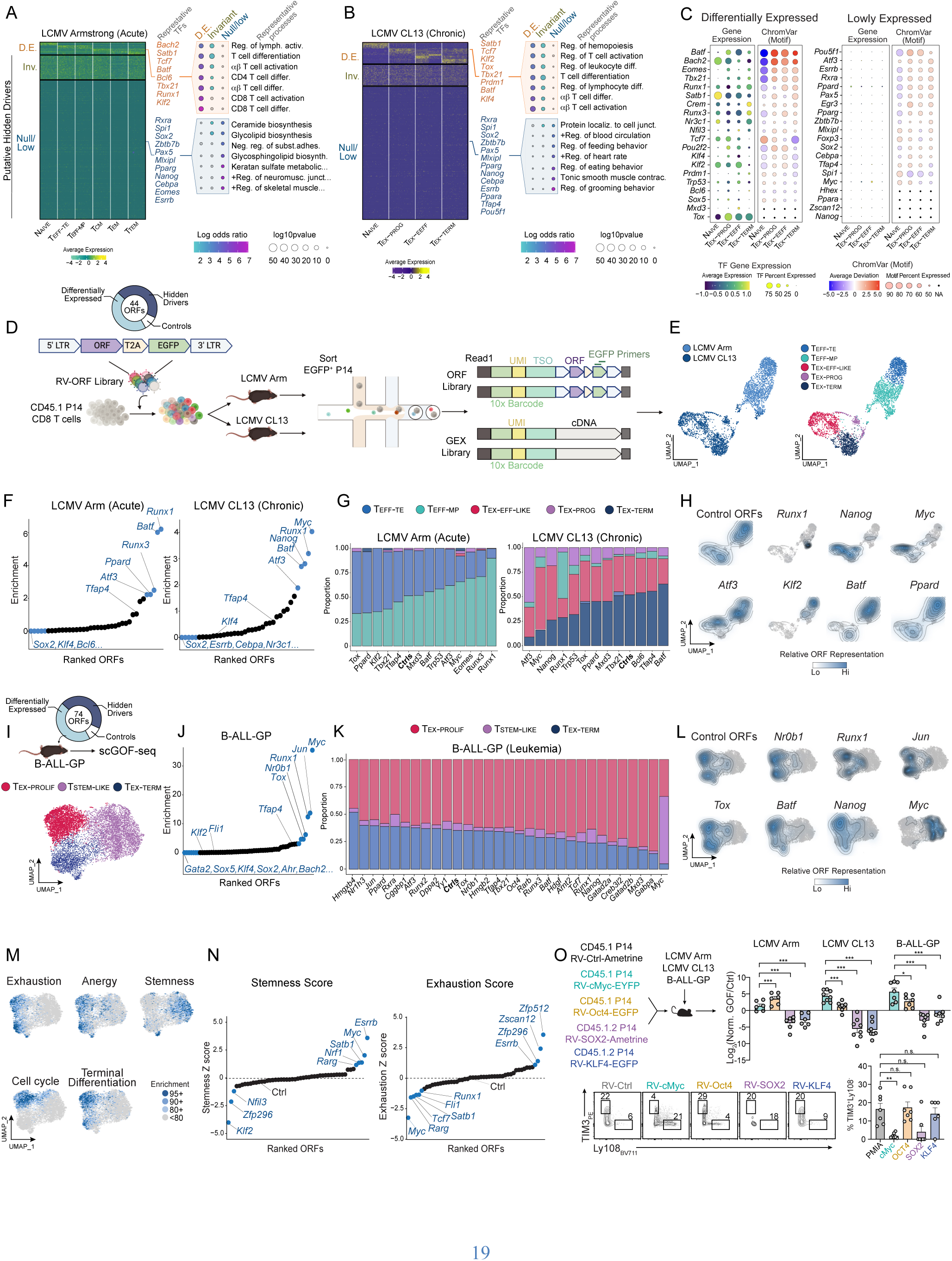
scGOF-seq screens uncover hidden regulators of CD8⁺ T cell differentiation *in vivo*. **(A)** Expression dynamics of differentially expressed (D.E.) genes, invariant (Inv.) genes, and lowly expressed genes in canonical CD8^+^ T cell states in LCMV Arm based on GSE290082 with representative TFs highlighted (left). Representative processes enriched based on gene ontology analyses of D.E. genes and lowly expressed genes (right). **(B)** Expression dynamics D.E., Inv., and lowly expressed genes in canonical CD8^+^ T cell states in LCMV CL13 from reanalysis of GSE290082 with representative TFs highlighted (left). Representative processes enriched based on gene ontology analyses of DE genes and lowly expressed genes (right). **(C)** Gene expression and motif deviation scores based on paired scRNA-seq and scATAC-seq profiling (GSE290082) of most TFs included in the initial screening library. TFs lacking known motifs are indicated as “NA.” **(D)** P14 CD8⁺ T cells were transduced with the RV-ORF library and transferred into mice infected with LCMV Arm or CL13. EGFP⁺ donor cells were sort-purified 2wks post-infection for 5’ scRNA-seq analysis. A paired ORF library was generated based on amplification of EGFP-containing transcripts to assign ORF identities. **(E)** scGOF-seq UMAP colored by condition and T cell states. **(F)** Representation of each ORF in LCMV Arm and LCMV CL13 normalized to input. **(G)** Composition of cell states for select ORFs (ORFs with sufficient recovery included). **(H)** Representative abundance of ORFs projected into the UMAP space from E. **(I)** P14 CD8⁺ T cells transduced with an expanded TF-GOF library (74 vectors) were transferred into mice bearing B-ALL-GP (top). EGFP⁺ donor cells were sorted 10 days post-transfer for scGOF-seq and projected in the UMAP space (bottom). **(J)** Representation of each ORF in the B-ALL-GP screen normalized to input. **(K)** Composition of cell states for each ORF from I **(L)** Representative abundance of ORFs projected into the UMAP space from I. **(M)** Gene module enrichment for exhaustion, anergy, stemness, cell cycle, and terminal differentiation signatures. **(N)** Gene module enrichment for stemness and exhaustion signatures for each ORF in B-ALL-GP scGOF-seq. **(O)** Mixed transfer analysis of cMyc GOF, Oct4 GOF, KLF4 GOF, SOX2 GOF, and Ctrl cells in LCMV Arm (D14), LCMV CL13 (D14), and B-ALL-GP (D12). Graphs show mean ± SEM of n=7 mice pooled from two independent experiments (Fig. 1O). *p<0.05, ***p < 0.001, n.s. not significant, paired Student’s t-test. For scORF-seq screens cells were pooled from n=5-11 mice (D) or n=2 mice (I).

To systematically interrogate GOF perturbations *in vivo*, we developed single-cell GOF sequencing (scGOF-seq), a pooled screening framework that links enforced ORF expression to transcriptional state at single-cell resolution in immunocompetent hosts. Notably, this platform enabled deconvolution of primary CD8^+^ T cells stably transduced with non-immunogenic, ORF-encoding vectors isolated from mouse tissues **(Fig. 1D)**. We generated a library of ∼45 vectors (including 4 ORF control vectors), each encoding ORF-T2A-EGFP cassettes composed of DE and putative hidden driver TFs (**Fig. 1C-D, fig. S2A**). Retrovirus was packaged in an arrayed format and subsequently used to transduce T cell receptor (TCR) transgenic P14 CD8⁺ T cells that recognize the GP_33-41_ epitope derived from Lymphocytic Choriomeningitis Virus (LCMV)(*40*). Transduced P14 cells were profiled for input transduction efficiency, mixed, and transferred to recipient mice subsequently infected with acutely resolving LCMV Armstrong (Arm) or persistent LCMV Clone 13 (CL13), well-established models for investigating T cell differentiation (**Fig. 1D**). Two weeks post-infection, transduced (EGFP⁺) P14 cells were sorted and processed for scRNA-seq, enabling ORF identity and gene expression profiles to be mapped to individual cells (**Fig. 1D; fig. S2B**). Through unsupervised clustering and gene-signature enrichment analysis (*5, 41*) **(fig. S2C)**, we resolved terminal effector cells (TEFF-TE), memory precursor effector T cells (TEFF-MP), terminally exhausted cells (TEX-TERM), progenitor exhausted cells (TEX-PROG), and effector-like exhausted cells (TEX-EFF-LIKE) (*38*) **(Fig. 1E)**.

As an initial validation, scGOF-seq accurately recapitulated anticipated regulatory logic. GOF of BATF and RUNX1 increased representation across both acute and chronic infection, while KLF2 GOF was selectively depleted under chronic antigen exposure, consistent with prior reports (*38, 42*) (**Fig. 1F**). State-resolved analysis further confirmed known associations, including enrichment of RUNX3 and RUNX1 ORFs in TEFF-MP cells(*11, 43, 44*), T-bet and KLF2 GOF in short-lived, TEFF-TE cells (*38*) and BATF GOF cells among TEX-TERM cells (*45*) (**Fig. 1G-H**). These results establish that scGOF-seq faithfully captures several established regulators of antiviral CD8^+^ T cell differentiation *in vivo*.

Beyond validation, scGOF-seq uncovered unanticipated regulatory capacity within the latent TF repertoire. GOF perturbation of several canonical stemness-associated TFs, including SOX2, OCT4 (encoded by *Pou5f1*), NANOG, KLF4, and cMyc, elicited striking and previously unrecognized effect on CD8⁺ T cell differentiation (**Fig. 1F-H**). SOX2, OCT4, and NANOG, which are minimally expressed across CD8⁺ T cell states, and cMyc, which is rapidly downregulated after activation, exhibited “hidden-driver” activity, as enforced expression induced differentiation outcomes absent in physiological settings. While KLF4 and SOX2 GOF broadly impaired T cell accumulation, cMyc and NANOG GOF selectively enhanced expansion during chronic infection (**Fig 1F**). Notably, scGOF-seq uncovered an unexpected role of cMyc GOF in constraining terminal exhaustion in LCMV CL13 (**Fig. 1G; fig. S2D**), which is in stark contrast to the well-established, native role of endogenous cMyc in driving terminal differentiation of CD8^+^ T cells(*33, 46–48*). In summary, scGOF-seq uncovered both anticipated and unanticipated patterns of representation of hidden driver and differentially expressed TF ORFs in LCMV infection settings.

### *In vivo* scGOF-seq uncovers synthetic TF programs with enhanced therapeutic potential in leukemia

GOF engineering holds direct therapeutic promise in the context of adoptive cell therapies in cancer. To assess whether these latent regulatory effects extend to cancer, we applied scGOF-seq to a syngeneic BCR-ABL *Arf^−/-^* B-cell acute lymphoblastic leukemia model expressing GP_33-41_ (B-ALL-GP) (*38*)(Fig. 1I), expanding the library to ∼75 ORFs (**Fig. 1I; fig. S3A-C**). Single-cell analysis of transduced P14 cells resolved TEX-TERM, proliferative exhausted cells (TEX-PROLIF), AND stem-like (TSTEM-LIKE) populations. As expected, established T cell regulators such as JUN, TOX, AP4, and RUNX1 enhanced T cell accumulation (*28, 42, 46, 49*) (**Fig. 1J**). Strikingly, the B-ALL scGOF-seq screen demonstrated cMyc GOF resulted in a ∼35-fold accumulation of P14 cells (**Fig. 1J**). The unbiased nature of scGOF-seq also demonstrated that cMyc GOF induced formation of a distinct population of tumor-specific cells characterized by reduced exhaustion and terminal differentiation signatures alongside increased stemness and persistence features (**Fig. 1K-N**, **fig. S3D**). Notably, these observed cell state-specific effects would be largely obscured in bulk genetic screens, highlighting the utility of scGOF-seq in uncovering unexpected regulators of T cell fate.

### scGOF-seq exposes latent functions of cMyc in programming T cell differentiation *in vivo*, distinct from other conventional pluripotency factors

In contrast to the well-established, native role of cMyc in driving terminal differentiation of CD8 T cells(*33, 46–48*), scGOF-seq revealed that cMyc GOF consistently promoted CD8⁺ T cell accumulation and stemness in both LCMV CL13 and B-ALL-GP **(Fig. 1F,J)**. To glean insights into these unexpected phenotypes driven by cMyc GOF, we leveraged cMyc-EGFP reporter mice to profile cMyc expression dynamics. Consistent with expression datasets implicating cMyc as a putative hidden driver in differentiated T cells **(Fig. 1C)**, endogenous cMyc expression was rapidly extinguished by day 6 *in vitro* and prior to day 10 *in vivo* in both infection and leukemia models **(fig. S4A-E)**, consistent with previous reports in LCMV Arm(*46*). Thus, the phenotypes observed with cMyc GOF may reflect activation of a latent regulatory program that is normally inaccessible during physiological T cell differentiation.

We next asked whether other canonical pluripotency-associated TFs were capable of inducing similar to phenotypes to cMyc GOF. Among stemness-related TFs, including OCT4, SOX2, KLF4, TCF7, GATA2, and others, only cMyc induced a *de novo* CD8⁺ T cell state enriched with features of self-renewal and effector competence under chronic antigen conditions **(Fig. 1I,M)**. Other pluripotency-associated TFs behaved fundamentally differently: OCT4 failed to enhance formation of TSTEM-LIKE populations **(Fig. 1K)**, while SOX2 and KLF4 GOF impaired T cell expansion **(Fig. 1F, J)**. To directly resolve these divergent outcomes, we performed head-to-head mixed-transfer analyses of cMyc, KLF4, SOX2, and OCT4 GOF perturbations across acute infection, chronic infection, and leukemia models **(Fig. 1O, fig. S4F)**. Under persistent antigen exposure, cMyc GOF drove >20-fold expansion, whereas KLF4 and SOX2 GOF resulted in 4 and 40-fold depletion, despite their shared roles as Yamanaka factors in non-immune lineages(*50*). Moreover, unlike cMyc, these factors failed to prevent terminal exhaustion *in vivo* **(Fig. 1O, fig. S4F-I)**. Collectively, these findings reveal previously unrecognized, context-dependent functions of cMyc and other pluripotency-associated TFs in T cells, underscoring the power of scGOF-seq to uncover hidden regulatory capacities that are unlikely to be predicted from endogenous expression patterns, LOF genetics, or bulk screening approaches.

### cMyc GOF overrides the terminal exhaustion program

Although scGOF-seq identified cMyc as a potent driver of CD8⁺ T cell accumulation, endogenous cMyc in native T cell contexts has been classically associated with terminal effector differentiation(*33, 46, 47*). Consistent with prior studies, cMyc-EGFP reporter P14 cells (*51, 52*) transferred into LCMV CL13-infected mice showed that cells with high levels of endogenous cMyc preferentially adopted terminally exhausted phenotypes **(fig. S5A)**. This behavior contrasted with the phenotype induced by enforced cMyc GOF, which instead promoted stem-like features and restrained exhaustion **(fig. S5B)**. To define the kinetics and context dependence of this effect, we co-transferred RV-Ctrl and RV-cMyc P14 cells into mice infected with LCMV Arm or CL13 strains and longitudinally profiled donor cells. Despite its canonical role in supporting CD8^+^ T cell proliferation, cMyc GOF transiently reduced P14 accumulation during early effector phases (days 5-8) in both models **(Fig. 2A)**. Strikingly, by day 15 and thereafter, cMyc GOF cells accumulated to a greater degree in chronic infection, reaching >20-fold enrichment by >day 42 in LCMV CL13 compared to only a 4-fold increase in acute infection. Thus, cMyc GOF confers context-specific, antigen-dependent accumulation.

**Fig. 2.**
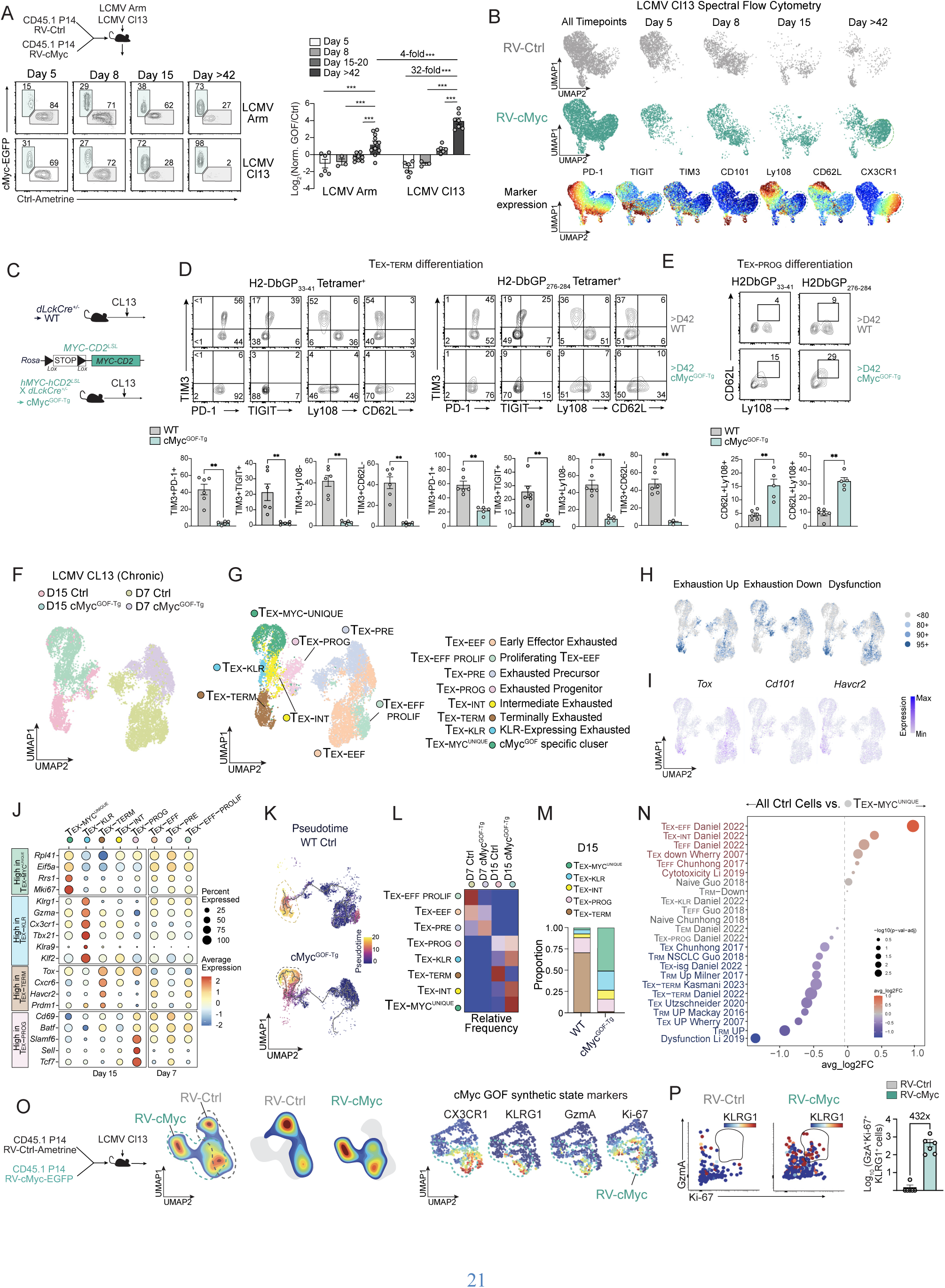
Artificially sustained cMyc expression restrains exhaustion differentiation. **(A-B)** RV-cMyc and RV-Ctrl P14 cells were mixed and transferred into congenic mice infected with LCMV CL13 or LCMV Arm. **(A)** Donor P14 cells enrichment were evaluated on days 5, 8, 15, and >42 post-infection by spectral flow cytometry. **(B)** UMAP representation and clustering of donor T cells from A (top) and relative expression of key proteins (bottom). Green dashed regions mark clusters predominantly composed of day 42 cMyc GOF cells. **(C)** WT and Myc^GOF-Tg^ mice were infected with LCMV CL13. Tetramer^+^ cells were analyzed via spectral flow cytometry >D42 post-infection. **(D)** The terminal exhaustion phenotype of H2-Db GP33–41 tetramer⁺ and H2-Db GP276–284 tetramer⁺ cells from panel C was analyzed based on co-expression of exhaustion markers (TIM-3, PD-1, TIGIT) and a TIM-3⁺ Ly108⁻ or CD62L⁻ profile, indicating loss of memory/progenitor markers. **(E)** TEX-PROG frequency via measuring CD62L^+^Ly108^+^ cells from C. **(F)** scRNA-seq analysis of cMyc GOF phenotype. CD8⁺ T cells were sort purified on days 7 and 15 post-infection from WT or Myc^GOF-Tg^ mice. UMAP representation and clustering of CD8⁺ T cells by condition. **(G)** UMAP representation and clustering of CD8⁺ T cells by cell state annotation according to key cell state-defining genes expression (fig. S6A-B, Fig. 2J). **(H-I)** Feature plots reflecting enrichment of exhaustion and dysfunction gene modules (H, top) and relative expression of exhaustion-associated genes, *Tox, Havcr2* (TIM3), and *Cd101* (I, bottom). **(J)** Gene expression analysis of key genes. (**K**) Trajectory and pseudotime analysis generated using Monocle3. Dashed regions indicate distinct terminal differentiation endpoints for WT and Myc^GOF-Tg^ cells. **(L)** Relative abundance of each cell state. **(M)** Summary of cell state composition of day 15 WT and cMyc^GOF-Tg^ mice based on scRNA-seq profiling. **(N)** Gene set comparison of WT control cells versus the TEX-MYC^UNIQUE^ population. **(O)** UMAP representation and clustering of RV-Ctrl and RV-cMyc P14 cells donor cells harvested from LCMV CL13-infected mice on day 14-16 of infection (right). Relative expression of GzmA, KLRG1, CX3CR1, and Ki-67 in donor cells from O (left). (**P**) Enumeration of GzmA^+^Ki67^+^KLRG1^+^ donor cells. Graphs show mean ± SEM of n=5-16 mice from one representative experiment or pooled from two or more independent experiments (Fig. 2A-G). *p < 0.05, **p < 0.005, **p < 0.001, paired Student’s t-test.

High-dimensional phenotyping revealed that cMyc GOF cells progressively diverged from control cells, displaying reduced expression of exhaustion markers (TIM3, TIGIT) and increased expression of stem-associated molecules (Ly108 and CD62L) **(Fig. 2B; fig. S5B-D)**. Notably, this divergence emerged only after day 15 of chronic infection, coincident with the onset of cMyc GOF-dependent T cell accumulation and after endogenous cMyc expression is extinguished **(fig. S4D)**. These temporal dynamics indicate that artificially sustained cMyc expression, outside its native regulatory window, may be required to unlock this synthetic differentiation program during chronic antigen exposure.

To determine whether cMyc GOF similarly blunted differentiation of endogenous, polyclonal exhausted CD8⁺ T cells, we generated cMyc GOF transgenic (cMyc^GOF-Tg^) P14 mice via crossing c*MYC-CD2^LSL^* mice (*53*) with distal Lck-Cre (*dLckCre^+/-^*) P14 mice, resulting in cMyc GOF in endogenous T cells (*54*) **(Fig. 2C)**. Although cMyc levels in transgenic cells were not as robust as those achieved in transduced cells **(fig. S5E)**, cMyc^GOF-Tg^ mice showed increased accumulation of GP33- and GP276-tetramer^+^ CD8⁺ T cells during LCMV CL13 infection **(fig. S5F)**. cMyc^GOF-Tg^ cells also exhibited reduced exhaustion phenotypes **(Fig. 2D)** and a greater frequency of Ly108⁺CD62L⁺ TEX-PROG cells with robust stemness features (*55*) **(Fig. 2E)**. Together, these results demonstrate that cMyc GOF limits exhaustion and preserves stem-like properties in both adoptively transferred and endogenous CD8⁺ T cells.

### Artificially sustained cMyc expression induces differentiation of a synthetic, effector-like state

To resolve how cMyc GOF reshapes exhaustion trajectories, we performed scRNA-seq on polyclonal CD8⁺ T cells isolated from cMyc^GOF-Tg^ and control mice on days 7 and 15 of LCMV CL13 infection **(Fig. 2F)**. Cells segregated by timepoint as expected, and annotation using established signatures and markers identified canonical exhausted states, including TEX-EEFF, TEX-EFF, TEX-PROG, TEX-INT, TEX-KLR, and TEX-TERM (*5*) **(Fig. 2G-J; fig. S6A-B)**. Trajectory analysis revealed a bifurcation at the TEX-INT stage: control cells progressed toward TEX-TERM, whereas cMyc GOF cells diverted into a distinct terminal branch, which we termed TEX-MYC^Unique^ **(Fig. 2G, K)**. Consistent with flow cytometry data **(Fig. 2B-E; fig. S5B-D)**, scRNA-seq analysis indicated that cMyc GOF markedly reduced formation of TEX-TERM cells, increased frequency of stem-like (TEX-PROG) and effector-like states (TEX-INT and TEX-KLR), and robustly expanded the TEX-MYC^Unique^ cell state **(Fig. 2L-M)**.

TEX-MYC^Unique^ cells exhibited reduced inhibitory receptor expression relative to TEX-TERM, but lower stem-associated gene expression than TEX-PROG **(Fig. 2J)**. Notably, this synthetic cell state displayed elevated *Cx3cr1* and *Klrg1* expression **(fig. S6B)** and were strongly enriched for effector-like transcriptional programs while being depleted of dysfunction signatures **(Fig. 2N; fig. S6C)**. However, despite sharing some effector-associated features, TEX-MYC^Unique^ cells were molecularly distinct from effector-like clusters (TEX-EFF states, TEX-EFF-PROLIF, TEX-INT and TEX-KLR; **Fig. 2G,J)**. A defining hallmark of TEX-MYC^Unique^ was a dramatic depletion of T cell dysfunction gene signatures **(Fig. 2N)** and marked upregulation of *Mki67* and ribosomal- and translation-associated genes such as *Rpl41*, *Eif5a*, and *Rrs1* **(Fig. 2J)**. Together, these findings support cMyc GOF drives formation of a discrete CD8 T cell state with proliferative effector qualities and elevated expression of genes associated with protein translation.

Consistent with the transcriptional features of the TEX-MYC^Unique^ cluster **(Fig. 2J; fig. S6B)**, spectral flow cytometry independently validated the emergence of this synthetic state **(Fig. 2O)**. cMyc GOF induced a *de novo* GzmA⁺Ki-67⁺KLRG1⁺ effector-like population in LCMV CL13 that was largely absent in control cells **(Fig. 2O)** and expanded at >400-fold higher frequency **(Fig. 2P)**. Because these experiments were performed in mixed-transfer settings, the appearance of this population likely reflects cell-intrinsic reprogramming rather than differences in viral burden or microenvironment.

Finally, tamoxifen-inducible activation of cMyc GOF in purified TEX-INT cells demonstrated a causal block in differentiation toward TEX-TERM, instead maintaining a CX3CR1⁺ effector-like cell state. In contrast, control CD8⁺ T cells followed an expected transition path from TEX-INT to TEX-TERM cells (*56*) **(fig. S6D-F)**. Collectively, these data show that cMyc GOF rewires exhausted CD8⁺ T cell differentiation, suppressing terminal exhaustion and generating a synthetic effector-like state that occupies a regulatory space inaccessible to natural T cell differentiation **(fig. S6G)**.

### cMyc GOF induces multimodal reprogramming of exhausted CD8⁺ T cells

cMyc coordinates expression of genes that support cellular fitness and stemness by promoting ribosome biogenesis, protein translation, glycolysis, lipid and amino acid metabolism, nucleic acid synthesis, mitochondrial biogenesis, proliferation, and DNA replication, among other pathways(*57, 58*). In contrast, metabolism, proliferation, and homeostasis are markedly altered in exhausted CD8⁺ T cells(*59, 60*). Although, cMyc is a classical regulator of CD8⁺ T cell activation, proliferation, and metabolism in early activated cells(*47*), cMyc is rapidly silenced following activation **(Fig. 3A; fig. S4A-E)**. Thus, cMyc GOF enforces sustained cMyc activity outside its physiological expression window in exhausted cells, constituting a non-physiologic, synthetic perturbation.

**Fig. 3.**
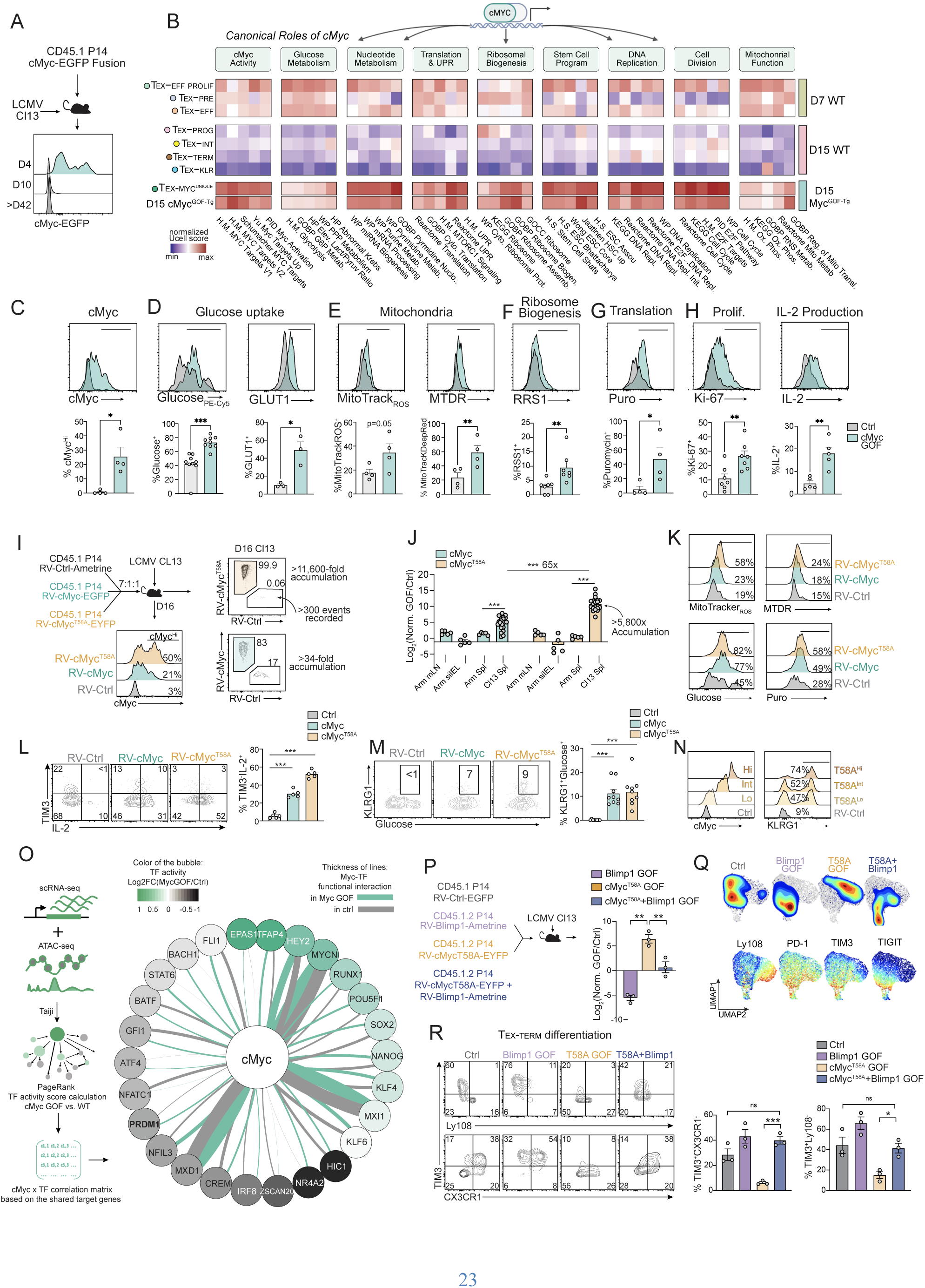
Multimodal reprogramming of exhausted T cells by artificially sustained cMyc activity. **(A)** cMyc reporter (cMyc-EGFP fusion) P14 cells were transferred into congenic distinct mice infected with LCMV CL13. Donor cells were analyzed on days 4, 10, and >42 post-infection by spectral flow cytometry. **(B)** Pathway enrichment analysis comparing canonical cMyc-associated programs across T cell states in WT cells and day15 MYC^GOF-Tg^ cells from the scRNAseq dataset in Fig. 2F. **(C–H)** Histograms and quantitative comparisons of cMyc expression, glucose uptake, GLUT1 expression, MitoTracker-ROS, MitoTracker Deep Red (MTDR), regulator of ribosome synthesis 1 (RRS1), puromycin uptake, Ki-67 expression, and IL-2 production in cMyc-RV versus control-RV cells in a mixed transfer setting in LCMV Cl13 on d12-16 of infection. **(I)** RV-Ctrl, RV-cMyc, and RV-cMyc^T58A^ P14 cells were mixed and transferred into mice infected with LCMV CL13. **(J)** Log2 accumulation of cMyc-RV and cMyc^T58A^-RV cells relative to RV-Ctrl cells in mLN, siIEL, and spleen during LCMV Arm or CL13 infection. Tissues were harvested on days 14–16 of infection. **(K)** Representative histograms comparing mitochondrial function, glucose uptake, and protein translation in donor cells from J. **(L)** TIM3 expression and IL-2 production on donor cells from J. **(M)** Glucose uptake and KLRG1 expression on donor cells from J **(N)** KLRG1 expression based on graded cMyc expression from J. **(O)** Taiji analysis of TF activity in cMyc-GOF cells. Color of circles indicates the TF activity difference between cMyc GOF vs. Ctrl. Line thickness represents the functional interaction or shared target genes between each TF and cMyc in cMyc GOF condition (green) and WT condition (gray). **(P)** RV-Ctrl, RV-BLIMP-1, RV-cMyc^T58A^, and RV-BLIMP-1+cMyc^T58A^ P14 cells were mixed and transferred to mice infected with LCMV CL13. Spleens were harvested on days 14–16 post-infection. **(Q)** Flow cytometry profiling of donor cells from P. **(R)** Representative flow cytometry plots and quantification of TEX-TERM differentiation, gated as TIM3⁺CX3CR1⁻ and TIM3⁺Ly108⁻ populations. Graphs show mean ± SEM of n=3-20 from one representative experiment or pooled from two or more independent experiments (Fig. 3A,C-N,P-Q). *p < 0.05, **p < 0.005, **p < 0.001, n.s. =non-significant, paired Student’s t-test, ***p < 0.001, **p < 0.01, *p < 0.05.

To define how cMyc GOF reshapes CD8⁺ T cell states, we analyzed canonical cMyc-regulated processes across effector, exhausted, and cMyc GOF cells using scRNA-seq of control (WT) and cMyc^GOF-Tg^ cells in the LCMV CL13 model **(Fig. 2F)**. In WT cells, cMyc-associated programs were broadly depleted in exhausted CD8⁺ T cells at day 15 relative to early effector-like states at day 7 **(Fig. 3B)**. In contrast, cMyc^GOF-Tg^ cells, including the TEX-MYC^Unique^ cell state, displayed a converse pattern, maintaining high enrichment of classical cMyc-driven pathways linked to cellular fitness, homeostasis, and stemness **(Fig. 3B)**.

We next examined how cMyc GOF rewires CD8⁺ T cells by experimentally validating cMyc GOF modulated metabolic pathways that support high biosynthetic activity during chronic antigen exposure **(Fig. 3C-G)**. Glucose uptake and surface expression of GLUT1 (*61*) were elevated in cMyc GOF cells **(Fig. 3D)**, indicating increased glucose utilization. MitoTracker assays showed increased mitochondrial mass and reactive oxygen species (ROS) production, reflective of enhanced respiratory activity and mitochondrial mass **(Fig. 3E)**(***62***). Together, these changes suggest that cMyc GOF may support the energetic and biosynthetic infrastructure required for sustained accumulation and function. Because metabolic and mitochondrial fitness are tightly linked to ribosome production and protein synthesis(*63–65*), we assessed anabolic output. cMyc GOF cells expressed higher levels of RRS1, a ribosome biogenesis factor also upregulated at the gene expression level **(Fig. 3F**; **Fig. 2J)** and showed an approximately sevenfold increase in nascent protein synthesis by puromycin incorporation (*66*) **(Fig. 3G)**. Since robust proliferation and effector function depend on extensive remodeling of translational and metabolic programs(*67, 68*), we assessed whether these changes translated into improved measures of proliferation and fitness. cMyc GOF cells indeed displayed increased Ki-67 expression and enhanced IL-2 production upon peptide restimulation, consistent with elevated proliferative and homeostatic capacity **(Fig. 3H)**. Together, these data show that enforced cMyc GOF synthetically sustains metabolic, mitochondrial, and translational programs in exhausted CD8⁺ T cells, accompanying enhanced accumulation and reduced exhaustion during chronic stimulation.

### Degradation-resistant cMyc further amplifies T cell expansion and metabolic reprogramming

cMyc protein abundance is tightly controlled by ubiquitin-mediated degradation, with phosphorylation at the T58 residue triggering rapid turnover(*69*). We therefore tested whether enforced expression of the stable cMyc^T58A^ isoform would further potentiate cMyc GOF phenotypes. In triple mixed transfer of RV-Ctrl, RV-cMyc, and RV-cMyc^T58A^ transduced P14 cells into LCMV CL13-infected mice, cMyc^T58A^ GOF cells exhibited substantially higher cMyc abundance than wild-type cMyc-expressing cells (**Fig. 3I**). While cMyc GOF increased P14 accumulation by 17-fold, cMyc^T58A^ GOF induced a dramatic >5,800-fold expansion of P14 cells, a 65-fold increase in the relative accumulation of cMyc^T58A^ cells compared with cMyc GOF in LCMV CL13. With cMyc^T58A^ GOF, some mice exhibited >11,000-fold enrichment of P14 cells within 16 days of infection, representing an unprecedented degree of accumulation of T cells in the widely used LCMV CL13 model (**Fig. 3I-J**).

At the phenotypic level, cMyc^T58A^ GOF broadly reinforced the metabolic and functional programs induced by wildtype cMyc GOF. cMyc^T58A^ GOF cells displayed equal or greater mitochondrial mass and activity, glucose uptake, translational output, IL-2 production, reduced inhibitory receptor expression, and increased KLRG1 compared with cMyc GOF cells (**Fig. 3K-M**), without altering IFNγ or TNFα production (**fig. S7A**). We also noted a dose-dependent relationship between cMyc abundance and key phenotypes, including KLRG1 expression levels (**Fig. 3N**).

Despite the oncogenic potential of stabilized cMyc, neither cMyc nor cMyc^T58A^ GOF induced unchecked or antigen-independent CD8⁺ T cell expansion. Instead, accumulation was context dependent, with only modest accumulation effects in acute LCMV Arm infection and no enrichment in non-lymphoid tissues such as the small-intestinal epithelium (**Figs. 1O, 2A, 3J**). To corroborate antigen dependence, we performed mixed transfers of cMyc GOF cells into irradiated hosts to assess antigen-independent, homeostatic expansion and transformation potential. Under these conditions, cMyc GOF cells failed to accumulate (**fig. S7B**). Moreover, cell-cycle checkpoint pathways remained intact based on gene-signature analyses (**fig. S7C**). Together, these data indicate that stabilized cMyc enhances antigen-driven T cell accumulation and metabolic fitness without inducing autonomous proliferation or overt transformation.

### cMyc reprogramming antagonizes Blimp1-dependent exhaustion programs

cMyc GOF led to multimodal changes in T cell biology, consistent with the broad-acting role of cMyc in transcriptional regulation. To elucidate mechanisms underlying cMyc-driven reprogramming of T cell exhaustion, we leveraged an integrated multiomic approach to infer TFs and their corresponding regulomes that are affected by cMyc GOF. We applied the Taiji framework, which integrates gene-expression and chromatin-accessibility data to predict TF activity(*70, 71*). Specifically, we combined ATAC-seq profiling of polyclonal CD8⁺ T cells isolated from WT and cMyc^GOF-Tg^ mice during LCMV CL13 infection and scRNA-seq data from matched mice to analyze TF activity and functional interactions of TFs with cMyc (**Fig. 3O**). Consistent with prior reports that cMyc potentiates AP4 activity(*46*), Taiji predicted elevated activity of AP4 (encoded by *Tfap4*) in cMyc GOF cells. In parallel, Blimp1 activity was predicted to be suppressed in the cMyc GOF context. Notably, we observed an increased overlap in predicted target genes between cMyc and Blimp1 in cMyc GOF cells compared to controls, suggesting functional antagonism at shared regulatory loci (**Fig. 3O**).

Blimp1 has been shown to counteract cMyc-driven phenotypes in B cells (*72–75*) and is a well-established driver of T cell exhaustion that restricts stemness programs(*76–79*), represses mitochondrial biogenesis(*80*), directly suppresses IL-2 production(*81*), and limits proliferation (*82*) under persistent antigen. We therefore hypothesized that cMyc GOF reprogramming operates, at least in part, through relief of Blimp1-mediated repression. To test this, we performed four-way mixed transfers of P14 cells transduced with RV-Ctrl, RV-Blimp1, RV-cMyc^T58A^, or RV-cMyc^T58A^+RV-Blimp1 into LCMV CL13-infected mice. Concomitant Blimp1 overexpression robustly abrogated the cMyc GOF-induced phenotypes, including enhanced T cell accumulation, stem-like features, effector programs, and reduced TIM3 expression (**Fig. 3P-R**). These data indicate that cMyc GOF reprograms exhausted CD8⁺ T cells in chronic infection in part by antagonizing Blimp1-dependent exhaustion programs, thereby enabling sustained metabolic activity, proliferation, and functional persistence.

### cMyc GOF reprograms T cell differentiation and enhances leukemia control

Having established that cMyc GOF drives sustained metabolic fitness, stem-like properties, and resistance to terminal exhaustion under chronic viral antigen exposure, we next investigated whether these same programs could be engaged in cancer. Leveraging B-ALL-GP scGOF-seq findings, we established that cMyc GOF increased P14 cell representation while concurrently enhancing stemness and reducing exhaustion signatures (**Fig. 1O**). Gene-set analyses revealed that cMyc GOF reshaped CD8⁺ T cell differentiation in B-ALL in a manner highly concordant with chronic LCMV CL13 infection, marked by enrichment of effector-associated programs and suppression of exhaustion and dysfunction signatures (**Figs. 2N and 4A**). Transcriptomic profiling data from B-ALL-GP scGOF-seq further demonstrated that cMyc GOF induced multimodal reprogramming (**Fig. 4B**), consistent with the reprogramming observed during chronic infection (**Fig. 3B**).

**Fig. 4.**
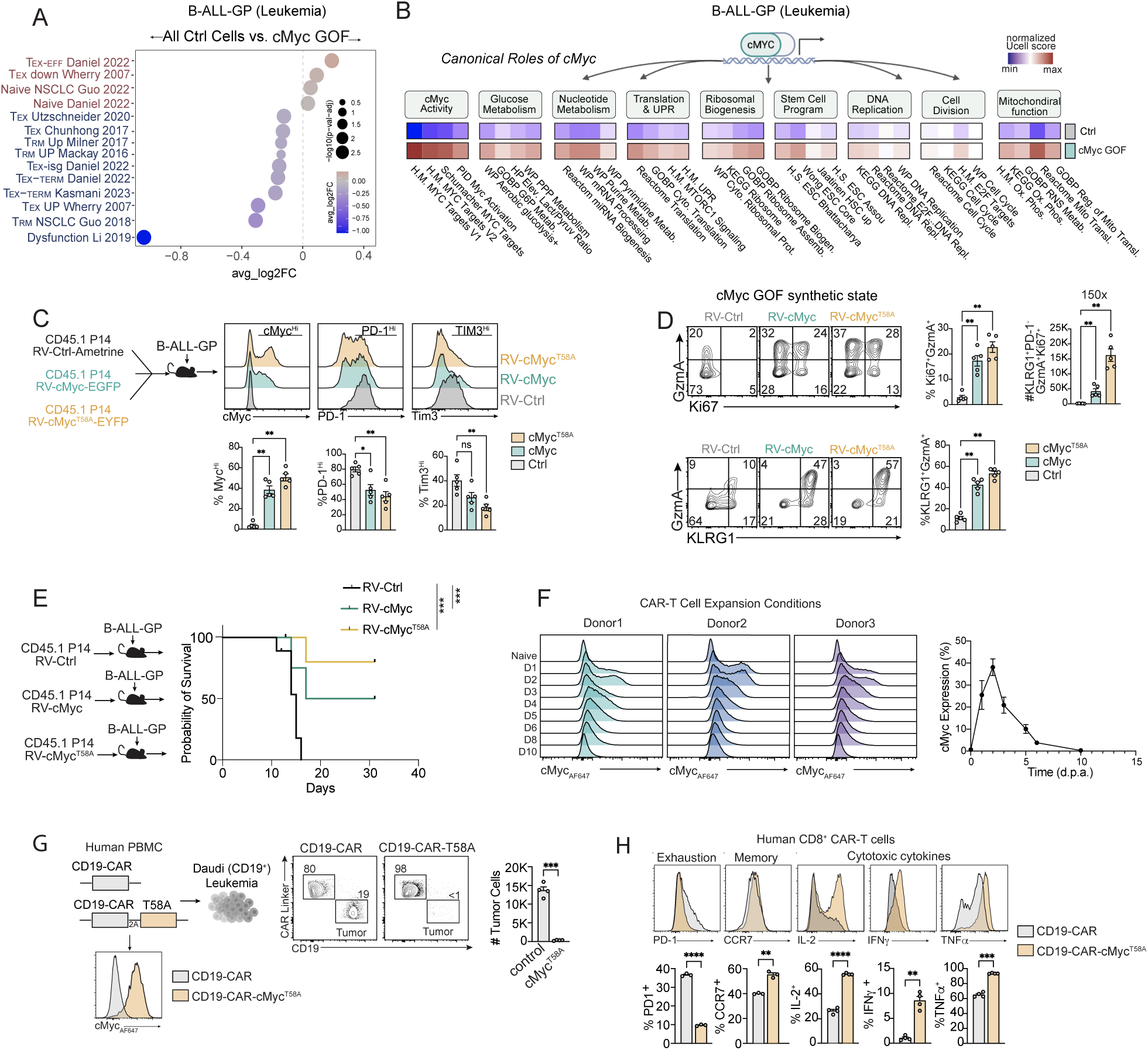
cMyc GOF enhances cell therapy efficacy in leukemia. **(A)** Gene set comparison of WT control cells versus the cMyc GOF population in B-ALL. **(B)** Pathway enrichment analysis comparing canonical cMyc-associated programs in WT control cells versus the cMyc GOF population in B-ALL. (**C**) Expression of cMyc, PD-1, and TIM3 in RV-Ctrl, RV-cMyc, and RV-cMyc^T58A^ transduced P14 donor cells transferred into B-ALL-GP bearing mice and analyzed on day 10 post T cell transfer. **(D)** cMyc GOF induced synthetic state identification of donor cells from C and enumeration of GzmA^+^Ki67^+^KLRG1^+^PD-1^−^ cells. **(E)** RV-Ctrl, RV-cMyc, or RV-cMyc^T58A^ transduced P14 were transferred individually into B-ALL-GP bearing mice, and survival was measured. **(F**) cMyc expression dynamics in human CD8⁺ T cells under standard CAR-T cell expansion conditions. **(G)** *In vitro* killing assay comparing human CD8^+^ T cells transduced with CD19-CAR (ctrl) and CD19-Myc^T58A^-CAR co-cultured with CD19^+^ Daudi Leukemia cells. **(H)** Expression levels of PD-1, CCR7, IL-2, IFN-γ, and TNFα in CD19-CAR and CD19-cMyc^T58A^-CAR T cells. Graphs show mean ± SEM of n=4-6 (Fig. 4C-E) or pooled from >3 independent donors (Fig. 4G-H). *p < 0.05, **p < 0.005, **p < 0.001, n.s.=non-significant, paired Student’s t-test, ***p < 0.001, **p < 0.01, *p < 0.05.

To validate these effects at the cellular level, we performed three-way mixed transfers of control, cMyc GOF, and cMyc^T58A^ GOF P14 cells into B-ALL-GP bearing mice. As in chronic infection, both cMyc and cMyc^T58A^ GOF cells maintained elevated cMyc expression and reduced inhibitory receptor levels (**Fig. 4C**), consistent with scGOF-seq results (**Fig. 1F-N**). Notably, cMyc GOF induced formation of a synthetic cell state (Ki-67⁺, GzmA⁺, PD-1^−^, and KLRG1⁺) similar to LCMV CL13 (**Fig. 2O**). In the B-ALL context, both cMyc and cMyc^T58A^ GOF cells exhibited increased frequencies of Ki-67⁺, GzmA⁺, and KLRG1⁺ cells, resulting in a ∼150-fold expansion of a GzmA⁺Ki-67⁺KLRG1⁺PD-1^−^ population that was absent among control cells (**Fig. 4D**).

We next asked whether these differentiation changes translated into therapeutic benefit. Adoptive transfer of cMyc GOF P14 cells significantly prolonged survival of B-ALL-GP bearing mice compared with controls (∼50% survival), while the stabilized cMyc^T58A^ isoform further improved survival to ∼80% (**Fig. 4E**). These data indicate that the cMyc-driven programs identified in chronic infection can be repurposed to enhance T cell efficacy in a leukemia setting.

Given the magnitude of these effects, we examined whether cMyc GOF similarly reprograms human CAR-T cells. Under standard CAR-T expansion conditions with IL-2, IL-7, and IL-15(*83*), activated human T cells exhibited transient cMyc induction (2–6 days post-activation) followed by rapid downregulation to near-baseline levels (**Fig. 4F**), paralleling observations in mouse T cells (**Fig. 4A and fig. S4FA-E**). We reasoned that this loss of cMyc activity during expansion may constrain CAR-T cell metabolic fitness and durability. To test this, we introduced cMyc^T58A^ into an anti-CD19-CD28 CAR construct. cMyc^T58A^-expressing CAR-T cells exhibited enhanced tumor cell killing (**Fig. 4G**), increased IL-2, IFN-γ, and TNF-α production, reduced PD-1 expression (**Fig. 4H**), and increased mitochondrial mass (**fig. S8A-B**), consistent with multimodal programming via cMyc GOF in mouse T cells. Together, these data indicate that sustained cMyc activity enhances the functional potency of human CAR-T cells and highlight cMyc GOF as a promising strategy for improving cellular therapies in hematologic malignancies.

### cMyc-induced stemness is insufficient for solid tumor control

Given that sustained cMyc expression enhanced ACT efficacy in B-ALL and promoted CD8⁺ T cell stemness, we next investigated whether cMyc GOF could similarly improve responses in solid tumors. We performed three-way mixed transfers of RV-Ctrl, RV-cMyc, and RV-cMyc^T58A^ P14 cells into mice bearing orthotopically implanted, syngeneic pancreatic ductal adenocarcinoma tumor cells (derived from C57BL/6 *Kras^G12D^Trp53^R172H^Pdx1-Cre* mice (*84*) stably expressing the LCMV GP_33-41_ peptide (KPC-GP) or mice bearing B16-GP tumors **(Fig. 5A-B)**. In both models, cMyc and cMyc^T58A^ GOF constrained terminal exhaustion and skewed differentiation towards Ly108⁺ TEX-PROG populations with cMyc^T58A^ GOF yielding a ∼2-4-fold increase relative to control. However, despite exhaustion prevention and a reinforced stem-like phenotype, cMyc^T58A^ GOF cells failed to control tumor growth or improve survival in B16-GP **(Fig. 5C)**. Thus, while cMyc GOF universally enhances stemness under persistent antigen exposure, cMyc^T58A^ GOF was insufficient to confer therapeutic benefit in solid tumors.

**Fig. 5.**
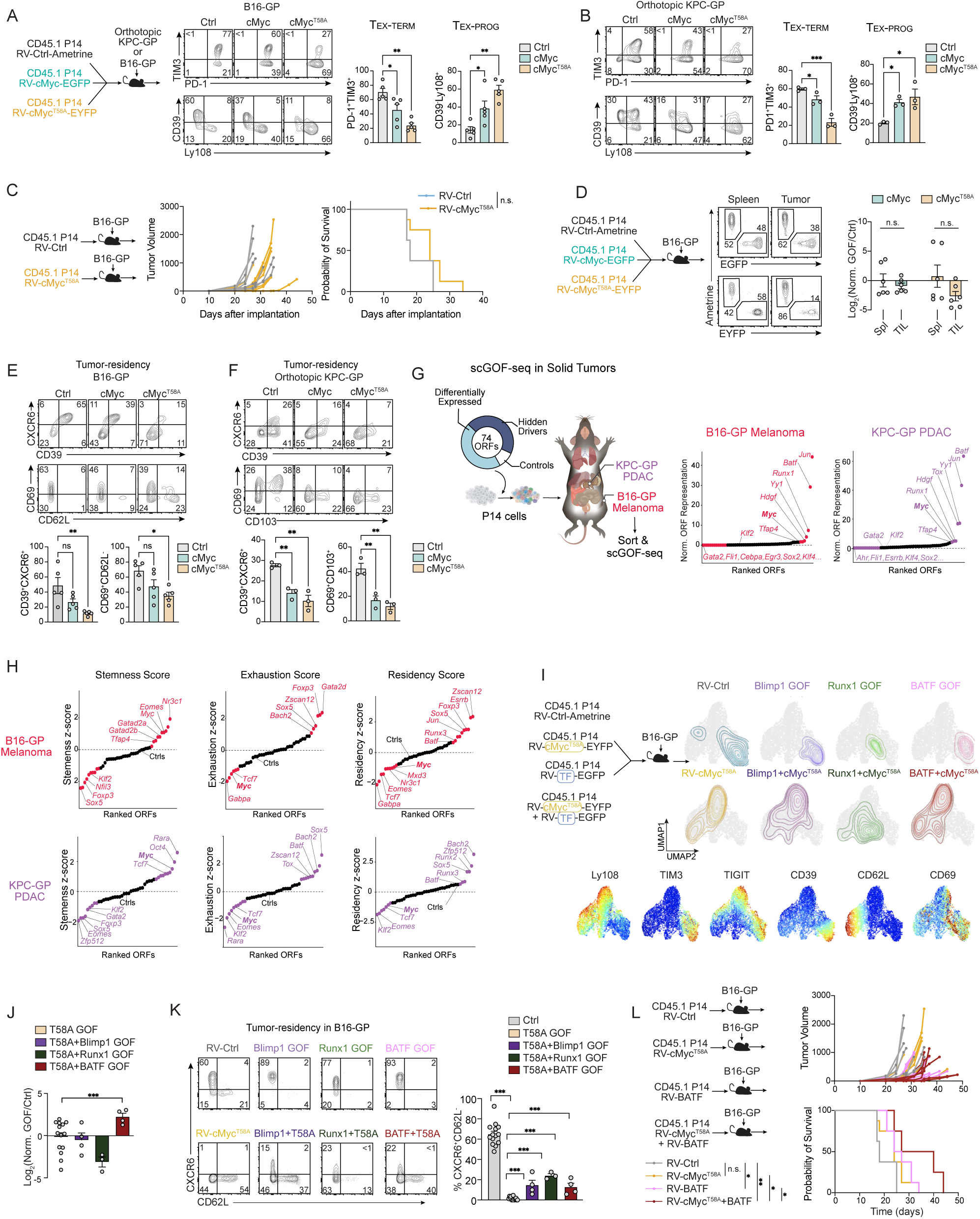
Rational GOF engineering enhances cell therapy efficacy in solid tumors. **(A-B)** Profiling of TEX-TERM and TEX-PROG markers of donor P14 cells transduced with RV-Ctrl, RV-cMyc, or RV-cMyc^T58A^ from mice bearing B16-GP (A) or KPC-GP tumors (B). T cells were harvested, isolated from tumors 8-12 days post-T cell transfer. **(C)** Tumor growth curves of B16-GP–bearing mice receiving RV-Ctrl or RV-cMyc^T58A^ transduced P14 cells. **(D)** Accumulation of RV-Ctrl, RV-cMyc, and RV-cMyc^T58A^ transduced P14 cells within B16-GP tumors shown in (A). **(E)** Tumor-residency phenotypes of RV-Ctrl, RV-cMyc, and RV-cMyc^T58A^ P14 cells from (A), assessed by co-expression of CD39 and CXCR6 or gating of CD69+ CD62L⁻. **(F)** Tumor-residency phenotypes of RV-Ctrl, RV-cMyc, and RV-cMyc^T58A^ P14 cells from KPC-GP tumors shown in (B), assessed by co-expression of CD39 and CXCR6 or CD69 and CD103. **(G)** scGOF-seq screening in B16-GP and orthotopic KPC-GP tumor models (left). Tumors were harvested 14 days after T-cell transfer, and normalized ORF representation was calculated for each model (right). **(H)** Gene-module enrichment scores for individual ORFs identified in (G). **(I)** Integrated analysis of combinatorial GOF perturbations assessed by spectral flow cytometry. Top, schematic of the *in vivo* combinatorial GOF strategy. CD45.1⁺ P14 cells transduced with RV-Ctrl, RV-cMyc^T58A^, single TF GOFs (Blimp1, RUNX1, or BATF), or dual GOFs combining each TF with cMyc^T58A^ were pooled and transferred into separate cohorts of B16-GP bearing mice. Cohorts included Ctrl + single cMyc^T58A^ GOF, Ctrl + single BLIMP1 + dual GOF, Ctrl + single RUNX1 + dual GOF, and Ctrl + single BATF + dual GOF. Donor P14 cells recovered from all cohorts from day 10-11 post-transfer were analyzed via spectral flow. Bottom, UMAP projections of recovered donor P14 cells, colored by GOF identity. Right, feature plots showing relative expression of differentiation and exhaustion markers (Ly108, TIM3, TIGIT, CD39, CD62L, CD69). **(J)** Accumulation of donor P14 cells from (I), normalized to input frequencies to enable quantitative comparison of enrichment across conditions. Tumor-resident populations were quantified by CD69⁺CD62L⁻ gating. **(L)** B16-GP–bearing mice received 2 × 10⁵ P14 cells transduced with RV-Ctrl, RV-cMyc^T58A^, RV-BATF, or RV-cMyc^T58A^ + RV-BATF, and tumor growth was monitored longitudinally. Graphs show mean ± SEM of n=3-14 (Fig. 5A-F) or pooled from >3 independent donors (Fig. 5K-M). *p < 0.05, **p < 0.005, **p < 0.001, n.s. =non-significant, paired Student’s t-test, ***p < 0.001, **p < 0.01, *p < 0.05. For scORF-seq screens in Fig. 5G, donor cells were pooled from n=8-10 mice.

Notably, cMyc GOF depleted tissue-residency programs **(Figs. 2N, 4A)** and failed to promote intestinal TRM state formation **(Fig. 3J)**. Tissue-residency programs are associated with enhanced CD8^+^ T cell adaption and retention in solid tumor environments(*38, 85*), we hypothesized that cMyc GOF failure in solid tumors is due to the accumulation within the tumor microenvironment (TME). Indeed, in both melanoma and PDAC, neither cMyc nor cMyc^T58A^ increased intratumoral or lymphoid accumulation **(Fig. 5D; fig. S9A).** Consistent with this finding, cMyc GOF reduced expression of residency markers (CD69, CD103, CXCR6) while increasing lymphoid-homing CD62L within tumors **(Fig. 5E-F)**. Multiomic analysis further suggested cMyc GOF limits activity of TFs supporting tissue-residency, including HIC1(*86*), Blimp1(*87, 88*), and BATF (*38*) **(Fig. 3O).** We therefore leveraged scGOF-seq to overcome the limited efficacy of cMyc GOF in solid tumors to identify TFs that could complement molecular programs and phenotypes driven by cMyc GOF in solid tumors **(Fig. 5G; fig. S9B-C)**. Screening 74 TF GOFs in B16-GP and KPC-GP tumors identified BATF and JUN as dominant drivers of P14 accumulation, whereas KLF2 reduced accumulation, consistent with prior work **(Fig. 5G)**(***12, 28, 38, 89***). scGOF-seq also captured expected roles for TCF1, RUNX3, and EOMES in regulating stemness and residency(*11, 38, 90, 91*), while uncovering unexplored regulators such as RUNX1 in enhancing TIL accumulation **(Fig. 5G)**. Informed by scGOF-seq, we also found that GATA2 blunts accumulation and features of exhaustion in tumor-specific T cells **(Fig. 5G; fig. S10A-B)**. Across both solid tumor models, scGOF-seq also demonstrated that cMyc GOF consistently promoted stemness while suppressing residency and accumulation, mirroring phenotypes observed in **Fig. 5A-F**.

### Combinatorial engineering integrates stemness and residency modules to drive superior efficacy in solid tumors

We next tested whether rational combinations of GOF perturbations could be leveraged to enhance cell therapy responses in solid tumors. Based on scGOF-seq results, we paired cMyc^T58A^ with BATF and RUNX1(TFs enhancing TIL accumulation from **Fig. 5G**) or BLIMP-1 (cMyc GOF antagonism identified from **Fig. 3O-R**) **(Fig. 5I)**. While all combinations partially restored residency-associated features, only BATF GOF increased intratumoral accumulation of cMyc^T58A^ GOF cells **(Fig. 5J)** and promoted CXCR6⁺ TRM -like states (*38, 92*) **(Fig. 5K)**. Functional testing revealed that although BATF GOF alone modestly improved tumor control(*12, 38, 89*), the cMyc^T58A^ + BATF combination produced the greatest tumor regression and survival benefit **(Fig. 5L)**. Collectively, these findings demonstrate that combinatorial GOF engineering, informed by scGOF-seq and exemplified by cMyc^T58A^ + BATF GOF, may be useful in supporting superior therapeutic outcomes in the context of cell therapies.

## Discussion

While LOF studies have extensively defined key regulators of T cell differentiation(*11–13, 70*), GOF perturbations have emerged as a complementary strategy for enhancing T cell activity(*20–23*). Stable TF overexpression imposes sustained and often “unnatural” expression kinetics and stoichiometries that can elicit synthetic activities distinct from native regulatory roles(*93*). Here, we performed antigen-specific scGOF-seq screens across five different immune competent models of disease. This high-resolution mapping demonstrated that enforcing TF activity beyond its physiological operating range can modulate differentiation pathways and, in some cases, induce the formation of T cell states not reported via native genetic regulation or conventional LOF genetics. Therefore, while GOF perturbations can amplify known biological processes, they can also induce *de novo* cell states that may be inaccessible through native signaling or LOF genetics.

The case of cMyc GOF exemplifies the principle of *de novo* state formation as a result of GOF engineering. Landmark studies showed that elevated endogenous cMyc in acute infection drives CD8⁺ T cells toward rapid effector commitment and terminal differentiation, and adoptive-transfer experiments demonstrated that cMyc^Hi^ cells exhibit limited self-renewal capacity(*33, 47, 94*). In line with these reports, we also found that CD8⁺ T cells with high levels of endogenous cMyc at early effector timepoints exhibit a more terminally differentiated phenotype. Thus, the discovery that artificially enforced cMyc expression enhances stemness was unexpectedly counter to its native role in T cells. Further analysis revealed a mechanistic basis for this divergence. Endogenous cMyc is rapidly downregulated following CD8⁺ T cell activation(*46*), and CD8⁺ T cells with relatively high endogenous *Myc* levels exhibited transcriptional profiles distinct from cMyc GOF cells associated with terminal exhaustion. These findings suggest that sustained endogenous cMyc expression may engage negative feedback circuits that actively constrain cMyc-driven stemness programs. We reasoned that enforced GOF bypasses these constraints, revealing a latent regulatory capacity of cMyc that was not accessible under physiological conditions. In contrast to the increased translation observed in exhausted TILs driven by GADD34-mediated p-eIF2α attenuation or ATF4 upregulation, which is associated with dysfunctional states(*95, 96*), the amplified anabolic programs induced by cMyc GOF instead enhanced proliferative fitness and cytokine production. The ability of cMyc GOF to counter exhaustion phenotypes was at least partly dependent on repression of Blimp1 activity, a TF known to limit mitochondrial fitness, IL-2 production(*97*), and T cell stemness(*78, 80*).

We also observed that cMyc GOF operates in a context-dependent manner. Despite its classical proto-oncogenic function(*98*), cMyc GOF did not result in ubiquitous, unchecked T cell expansion as might be anticipated of transformed cells. For example, we noted that cMyc GOF blunted T cell accumulation during the first week of acute or chronic infection. Further, cMyc GOF failed to enhance accumulation in non-lymphoid tissues or solid tumors. The expansive accumulation conferred by cMyc GOF in infection was predominantly antigen-dependent, as the accumulation of cMyc GOF cells in acutely resolved infection was modest compared with persistent infection and we did not detect accumulation of donor P14 cells in the absence of cognate antigen. Beyond these findings suggesting that cMyc GOF did not induce cellular transformation of mature CD8^+^ T cells, contemporary adoptive T cell therapies increasingly incorporate safety mechanisms such as caspase-inducible kill switches or other conditional control strategies to enable rapid elimination or regulation of transferred cells (*99–103*). Together, these observations indicate that cMyc GOF-based T cell engineering can be implemented within established safety frameworks, mitigating concerns associated with GOF perturbation of TFs. In this context, multimodal reprogramming, such as that mediated by cMyc or cMyc^T58A^ GOF, may be necessary to overcome the multitude of immunosuppressive hurdles encountered by T cells in cancer. Accordingly, our data highlight cMyc GOF as a promising approach to augment T cell responses in cancer. This notion corresponds with recent work highlighting that expression of certain oncogenes can enhance preclinical CAR-T cell responses(*27*).

Finally, the ability to strategically combine GOF perturbations, informed by approaches such as scGOF-seq, may expand the landscape of programmable T cell states with therapeutic potential. We identified a cooperative genetic interaction between cMyc and BATF GOF that illustrates how enforced co-activity can reprogram differentiation circuits beyond what single-factor perturbations achieve, suggesting that higher-order combinations may yield increasingly potent or durable T cell states. These findings are in line with recent efforts identifying the downstream cMyc effector AP4 as a regulator of T cell fitness that effectively pairs with BATF GOF(*24*). In parallel, rational integration of GOF and LOF perturbations(*25*), which can be guided by recent multi-omics-guided TF discovery pipelines(*70*), represents a promising approach to reprogramming T cell fate in cancer. Integrating these resources with scGOF-seq and extending them through AI/ML models trained on TF co-expression, network topology, and chromatin-inferred accessibility could enable predictive design of minimal TF “recipes” tailored to specific disease contexts.

To date, most immunotherapy strategies have focused on releasing inhibitory brakes, such as blocking PD-1 signaling or suppressing exhaustion-inducing TFs, whereas our work provides a roadmap for actively harnessing latent cell-state programs through GOF activation of hidden regulatory drivers. Together, technical and conceptual frameworks explored herein may provide a foundation for computation-guided engineering of synthetic immune programs that enable tailoring of cell therapy approaches for improved efficacy.

## Acknowledgments

The authors are extremely grateful for the assistance of Carlton Anderson and Gabrielle Cannon from the UNC Advanced Analytics Core (UNC Center for GI Biology and Disease, P30 DK034987) for assistance with FACS and scRNA-Seq library preparation. We further acknowledge the technical support from the UNC High Throughput Sequencing Facility. This facility is supported by the University Cancer Research Fund, Comprehensive Cancer Center Core Support grant (P30-CA016086) and the UNC Center for Mental Health and Susceptibility grant (P30-ES010126).

## Author contributions

Conceptualization: BMP, GNM, HKC, JJM

Data curation: BMP, GNM, HKC, JJM

Formal analysis: BMP, GNM, NB, FX, JM, VZ, AJ, HKC, JJM

Investigation: BMP, GNM, WDG, FX, LVR, JG, EFM, HKC, JJM

Methodology: BMP, GNM, JM, HKC, JJM

Resources: TE, GD, HKC, JJM

Supervision: WW, AS, JET, HKC, JJM

Visualization: BMP, GNM, NB, JM, VZ, HKC, JJM

Funding acquisition: HKC, JJM

Project administration: HKC, JJM

Writing – original draft: BMP, GNM, HKC, JJM

Writing – review & editing: BMP, GNM, HKC, JJM

## Data, code, and materials availability

No original code was created in this manuscript. All packages and scripts were modified from publicly available sources for case specific uses. All data and code are available upon request. All sequencing datasets generated for this study have been deposited at GEO: GSE302609, GSE302580, and GSE309695 (scGOF-seq) and will be made publicly available as of the date of publication. Information and requests for reagents, code, and data should be directed to the lead contacts, H. Kay Chung or Justin Milner. All materials will be made available upon request.

## Supplementary Materials

Materials and Methods

Supplementary Text

Figs. S1 to S10

Tables S1 to S3

Data S1 to S3

