## Supplemental Material for "Latent Regulatory Programs Generate Synthetic T Cell States with Enhanced Therapeutic Potential"

#### Materials and Methods

Mice

All mice were bred and housed in specific pathogen-free conditions on a 12-hr light-dark cycle at ambient temperature in accordance with Institutional Animal Care and Use Guidelines of the University of North Carolina, Chapel Hill. CD45.1 or CD45.2 C57BL/6 mice aged between 6 and 18 weeks old were used as recipients for cell transfer experiments. Transgenic TCR (P14) mice that recognize the LCMV peptide $GP_{33-41}$ were gifted by Dr. Ananda Goldrath (UCSD). P14 $cMyc^{GOF\text{-}Tg}$ mice were generated by crossing dLck-Cre (*44*) P14 mice to *Gt(ROSA)26 Sor*$^{tm13(CAG\text{-}MYC,\text{-}CD2*)Rsky}$/J mice (*43*) obtained from the Jackson Laboratory (JAX strain 033805). P14 ERT2-Cre h*MYC*-h*CD2*$^{LSL}$ (iCre $cMyc^{GOF\text{-}Tg}$) mice were generated by crossing ERT2-Cre P14 mice with *Gt(ROSA)26 Sor*$^{tm13(CAG\text{-}MYC,\text{-}CD2*)Rsky}$/J mice. Mice with EGFP fused to the 3' end of cMyc exon 3 (*88*) ($Myc^{tm1.1Dlev}$/J, strain 019075) were obtained from JAX, then crossed to P14 transgenic mice. Mice were bred in-house and fed standard Purina chow. Donor and recipient mice were either sex-matched or female cells were transferred into male mice. Both male and female mice were used.

T cell transfers, LCMV infection, and treatments

For adoptive transfer of T cells, 2.5-5.0 x $10^3$ naïve P14 cells were transferred into congenially distinct recipient mice one day prior to intravenous infection with 2-4x$10^6$ PFU LCMV CL13 or 2-3x$10^4$ naive P14 cells were transferred one day prior to intraperitoneal infection with 2x$10^5$ PFU LCMV Arm.

For adoptive transfer of transduced T cells, negatively enriched $CD8^+$ T cells were first isolated using the EasySep Mouse $CD8^+$ T cell Isolation Kit (StemCell, cat# 19853A) following manufacturer's instructions, then activated in wells coated with 100 µg/mL goat anti-hamster IgG (H+L; Invitrogen), 1 µg/mL plate-bound anti-CD3 (145-2C11; eBioscience), and 1 µg/mL anti-CD28 (37.51; eBioscience). Approximately 18-24h after activation, CD8 T cells were transduced with retroviral vectors as described previously (*29*). One day after transduction, transduced T cells were mixed, input ratio was measured, and 1-5 x $10^5$ cells were transferred into congenic mice infected with LCMV. Retroviral vectors used for these studies are outlined in Table S1.

For sort transfer experiments, $PD\text{-}1^+CX3CR1^+Ly108^-KLRG1^-$ $CD8^+$ T cells were sorted from LCMV CL13 infected iCre $cMyc^{GOF\text{-}Tg}$ mice 20 days post-infection, and 5 x$10^4$ donor cells were transferred to infection matched recipient mice. Recipient mice were treated with 200 µg of anti-CD4 (GK1.5; BioXcell) the day before and day following LCMV CL13 infection to deplete CD4 T cells for deeper T cell exhaustion. Tamoxifen-induced activation of cMyc GOF was achieved by intraperitoneal injection of 16.7 µg/mL tamoxifen diluted in sunflower oil 24 and 48 hours post transfer of sorted cells. Transferred CD45.2 iCre $cMyc^{GOF\text{-}Tg}$ cells were profiled 12 days post transfer via spectral flow cytometry.

Tumor models

All cell lines were maintained in sterile culture conditions at 37°C and 5% $CO_2$, at or below 80% confluency, and were routinely tested for Mycoplasma. All tumor cell lines (B16-F10 melanoma (obtained from Dr. Ananda Goldrath), KPC-4662 pancreatic ductal adenocarcinoma (obtained from Dr. Robert Vonderheide), and B-ALL leukemia (obtained from Dr. Martine Roussel) used in this paper express the LCMV $GP_{33-41}$ peptide and were previously generated and validated (*29*). B16-GP and KPC-GP cell lines were grown in complete DMEM with 10% FBS. The B16-GP growth media was supplemented with 300 μg/mL of geneticin. B-ALL-GP cells and Daudi cells (obtained from Dr. Gianpietro Dotti (UNC-CH)) were grown in complete RPMI with 10% FBS.

For B16-GP experiments, 0.5-1 x $10^6$ B16-GP cells were implanted subcutaneously into the flank 7-10 days before adoptive transfer of 1-2 x $10^6$ activated and expanded P14 T cells. P14 T cells were expanded with 20 U/mL rIL-2 (Peprotech), 2.5 ng/mL rIL-7 (Peprotech), and 2.5 ng/mL rIL-15 (Peprotech) for 2-4 days prior to transfer to congenic, tumor-bearing mice. For P14 cell efficacy experiments, 7.5 x $10^5$ B16-GP cells were implanted and 2.5 x $10^5$ transduced P14 T cells were transferred on the same day of transduction to maximize engraftment. Mice were euthanized if tumors ulcerated or reached 20mm in any direction. For B-ALL-GP phenotyping experiments, mice were inoculated intraperitoneally with 1.5 x $10^4$ B-ALL-GP cells one day before intravenous transfer of 2 x $10^3$ activated and transduced P14 T cells. For B-ALL-GP efficacy experiments, 5 x $10^3$ B-ALL-GP cells were implanted and 2 x $10^4$ transduced P14 T cells were transferred the following day. For experiments with KPC-GP tumor-bearing mice, 1.5-2.5 x $10^5$ KPC-GP cells were surgically implanted orthotopically into the tail of the pancreas as described previously (*29*). 7-10 days after KPC-GP implantation, 1-2 x $10^6$ transduced P14 T cells were adoptively transferred.

Tissue processing, spectral flow cytometry, and cell sorting

Single-cell suspensions from spleens were obtained through mechanical disruption and RBCs were lysed with ACK buffer (150 mM NH4Cl, 1mM $KHCO_3$, and 0.1 mM EDTA, pH 7.4). Intraepithelial lymphocytes (IELs) were obtained from small intestines through chemical digestion in a buffer containing 154 μg/mL DTE for 30 min at 37°C following removal of Peyer's patches. Tumor infiltrating lymphocytes were obtained from tumors through mechanical disruption and chemical digestion in RPMI with either 100 IU/mL Collagenase I (Worthington), 1mM MgCl2, 1mM CaCl2, 1% HEPES, 1 mM L-glutamine, and 5% bovine growth serum for B16 tumors or 1.25 mg/mL Collagenase IV (Worthington), 1 mg/mL DNAse I (Worthington), 0.1% Trypsin Inhibitor from Soybean (Worthington), and 5% bovine growth serum for KPC tumors for 30 min at 37°C. Following digestion, single-cell suspensions of lymphocytes were obtained from tumor and small intestines by filtering cells through a 70-μm nylon filter and then density gradient spin isolation was performed using 44/67% Cytiva Percoll Centrifugation Media (Fisher Scientific, cat# 45-001-747) for 20 min at 568g.

For flow cytometry analysis, all samples were stained with fixable live/dead dye (Invitrogen) and FcR block (S17011E; Biolegend) prior to further staining. The H-$2D^b$-$GP_{33-41}$ and H-$2D^b$-$GP_{276-286}$ tetramers were obtained from the National Institutes of Health Tetramer Core. The following antibodies were used in flow staining and purchased from Biolegend, unless otherwise noted: CD8α (53-6.7), CD8β (YTS156.7.7), CD45.1 (A20), CD45.2 (104), Tim3 (RMT3-23), Ly108 (330-AJ and 133G3; BD Biosciences), PD-1 (29F.1A12), TIGIT (1G9), CX3CR1 (SA011F11), CD62L (MEL-14), CD103 (2E7), CD69 (H1.2F3), CD127 (A7R34), CD44 (IM7), anti-human CD2 (TS1/8), Lag3 (C9B7W), CXCR6 (SA051D1), CD39 (Duha59), KLRG1 (2F1; Invitrogen), CD101 (Moushi101; Invitrogen), puromycin (2A4), IL-2 (JES6-5H4), IFN-γ (XMG1.2), TNF (MP6-XT22), GzmB (QA16A02), GzmA (3G8.5), BATF (S39-1060; BD Biosciences), TOX (REA473; Miltenyi Biotec), Ki67 (SoLA15; Invitrogen), TCF1 (C63D9; Cell Signaling), cMyc (E5Q6W; Cell Signaling), and anti-rabbit IgG (polyclonal; Cell Signaling). Intracellular staining was performed with the FoxP3 Transcription Factor Staining Kit (Invitrogen) according to manufacturer's instructions following 5-8min fixation with 2% PFA at room temperature.

For metabolic assessment of CD8⁺ T cells by flow, cells were treated with uptake and mitochondrial dyes as described previously (*89*). In brief, cells were incubated with 15 nM Mitotracker Deep Red (ThermoFisher) and 100 nM Mitotracker Red CMXRos (ThermoFisher) for 10 min at 37°C. Cells were then washed in cold PBS prior to extracellular staining. Alternatively, cells were incubated with 0.4 µM glucoseCy5 (Sigma Aldritch) for 30 min at 37°C. Cells were then washed in cold PBS before extracellular staining. Protein translation was assessed as similarly described in the SCENITH assay (*56*), where Puromycin was introduced into media of ex vivo cultured $CD8^+$ T cells for 30 min prior to staining of cells and intracellular staining of puromycin followed by spectral flow cytometry analysis.

All flow cytometry analysis was performed on a 5L Cytek Aurora spectral flow cytometer. All cell sorting was performed on a BD Aria II or Sony SH800 sorter. For LCMV sorts, CD4 T cells, RBCs, B cells were depleted from cell preparations on Miltenyi Biotec magnetic columns using biotin conjugated antibodies, CD4 (GK1.5), CD45R/B220 (RA3-6B2), CD19 (8D5), and TER-119 (TER-119), and streptavidin conjugated magnetic beads prior to sorting.

<u>Human CAR-T cell studies</u>

*Human CAR-T cell preparation.* Human CAR-T cells were generated from peripheral blood mononuclear cells (PBMCs) isolated from fresh human blood (Gulf Coast Regional Blood Center). PBMCs were resuspended in complete CAR-T medium (1:1 RPMI 1640: Click's medium supplemented with 10% FBS, GlutaMAX, penicillin/streptomycin, and 0.05 mM 2-mercaptoethanol) and activated on non–tissue culture–treated 24-well plates pre-coated with anti-human CD3 (Cytek) and anti-human CD28 (BD Pharmingen) monoclonal antibodies (1 µg/mL each). One day after activation, IL-7 (10 ng/mL) and IL-15 (5 ng/mL) were added. On day 2 post-activation, T cells were transduced with either control CD19.CAR or cMycT58A.CD19 CAR retroviral supernatant on retronectin-coated 24-well plates: wells were coated overnight with retronectin (7 µg/mL), loaded with viral supernatant, and centrifuged at 2,000×g for 90 min at 32°C before adding 5e5 activated T cells per well in cytokine-containing CAR-T medium, followed by centrifugation at 1,000×g for 10 min at 32°C. Cells were then cultured at 0.5-1 x $10^6$ cells/mL in IL-7/IL-15 supplemented CAR-T cell media, with media and cytokines refreshed every 2–3 days. CAR-T cells were used for experiments between days 8 and 12 post-activation.

*In vitro cancer killing assay.* Co-culture of CAR T cells and Daudi cancer cells was performed as previously described (*12, 90*). Daudi cells were seeded in 96-well plates at 1e5 cells/well and co-cultured at a 1:1 ratio with either control CD19.CAR T cells or cMycT58A.CD19 CAR T cells. Cells were cultured for 6 days in the presence of 100 IU/mL rhIL-2 before assessment of T cell exhaustion and cytokine production by flow cytometry.

*Mitochondria and ER confocal microscopy of human CAR-T cells.* CD8+ CAR-T cells were sorted on viable CD8+/GS4 (CAR marker)+ gates. Cells were blocked with 2% BSA in PBS for 1 hour at room temperature before antibody staining with rabbit α-Sec61b (Cell Signaling) and mouse α-Tom20 (Santa Cruz) antibodies in 1% BSA in PBS for 1 hour at room temperature. Cells were washed with 1x PBS and incubated with nuclear stain DRAQ5 (Abcam), and secondary antibodies goat anti-rabbit IgG (H+L) AlexaFluor 594 and goat anti-mouse IgG (H+L) AlexaFluor 488 (ThermoFisher) in 1% BSA in PBS for 1 hour at room temperature. A 96-well glass bottom plate (Cellvis) was coated with CellTak (Corning), washed with DI water, air dried, and labeled cells were centrifuged for adhesion. Cells were imaged using the Zeiss LSM-880 confocal microscope with ZEN acquisition software (Zeiss) or AiryScan detector and 3-D reconstruction was generated from confocal z-stacks using Imaris software. All quantitation indicates a per-cell measurement (i.e., a single point in the Mitochondria area/Nucleus area plot refers to the area of positive TOM20 signals above a manually-defined threshold, divided within a single cell, divided by the area of the nucleus (DRAQ5 positive signal from a manually-defined threshold) for that same cell. The threshold for both TOM20 and DRAQ5 signals was identical for segmentation for both samples. Manually-identified cells with irregular morphology or appearance representative of an apoptotic cell were excluded from calculations.

Retroviral Transductions

All mouse $CD8^+$ T cell transductions were performed as described previously (*91*) using packaged MSCV retrovirus, supplemented with 8 µg/mL polybrene and 50 µM BME, via spin-fection for 1 hr at 37°C and 568g. In addition to vectors included in the ORF screen (Table S1), the following vectors were obtained from Vector Builder, Addgene, or subcloned with the In-Fusion cloning approach: cMyc-IRES-EYFP (Vector Builder), $cMyc^{T58A}$-IRES-EGFP (Addgene; cat#177648), $cMyc^{T58A}$-IRES-EYFP (Vector Builder), and IRES-Ametrine (Vector Builder). Retrovirus was generated as previously described (*91*). In brief, PLAT-E retrovirus packaging cells were treated with TransIT-LT1 transfection reagent mixed with 1.5 µg of vector plasmid and 1 µg pcl-eco helper plasmid (1.5:1 ratio) at a 3:1 ratio of TransIT-LT1to DNA. Retrovirus was harvested 48 and 72h following transfection.

scGOF-seq Library Preparation and Sequencing

*scGOF-seq screening library and sample preparation.* All plasmids for the ORF screen were obtained from VectorBuilder on an MSCV backbone with an EGFP reporter. Enriched and activated P14 $CD8^+$ T cells were transduced with retrovirus in an arrayed format. For LCMV screens, 3e5 pooled cells were transferred into recipient mice one day after infection with LCMV Arm or LCMV CL13. On day 14 of infection $EGFP^+$ donor cells were sort purified. For B-ALL-GP screening, 2.5e5 pooled P14 T cells were transferred to recipient mice inoculated with 1.5e4 B-ALL-GP cells one day prior. For B16-GP screening, 1e6 pooled P14 T cells were transferred into mice bearing B16-GP cells implanted 10 days prior to T cell transfer. For KPC-GP samples, 2e6 pooled cells were transferred into mice bearing orthotopically implanted KPC-GP cells 10 days prior to T cell transfer. For all screens, $EGFP^+$ donor $CD8^+$ T cells were sorted from either splenocytes (LCMV Arm, LCMV CL13, B-ALL) or tumor (B16, KPC) on a Sony SH800 sorter. B-ALL-GP samples were sorted 10 days after T cell transfer, B16-GP samples and KPC-GP samples were sorted 14 days after T cell transfer.

*Processed scGOF-seq data analysis.* Cells from each sample were processed independently via the Seurat package (v5.1.0) in R (v4.3.3). Data were filtered to exclude any cells with more than 12% of reads derived from mitochondrial sequences, fewer than 3000 UMI counts, and fewer than 300 genes. Data were normalized via standard LogNormalization, and 3000 variable features were used for PCA dimension reduction. The first 30 principal components were used as input for neighbor graph construction and UMAP dimension reduction. Clustering was performed at multiple resolutions via the Louvain algorithm with multi-level refinement.

Input cell proportions were used to calculate ORF enrichment by normalizing ORF frequencies in experimental samples to their corresponding ORF and control proportions in the input donor cell population. ORF outputs normalized by input are available in **Data S1**. Enrichment scores were ranked and visualized accordingly. To generate ORF density plots, we used Nebulosa (*92*) to recover and smooth signals from sparse features in single-cell datasets. gRNA density plots were generated using scCustomize (*93*), which applies kernel density estimation to visualize guide RNA distributions across cell states. Gene signature log2 foldchange comparison between Ctrl vs cMyc GOF in Fig. 4A was performed using the R package escape (2.6.1, (*94*)). Gene signature and metabolic pathway enrichment were quantified using the R package UCell (v2.2), which computes enrichment scores based on the Mann–Whitney U statistic (*95*). UCell scores for each ORF were Z-score normalized and visualized as ranked enrichment profiles. Summary of scGOF-seq hits is available in **Data S2**. Gene sets used in this study is available in **Data S3**.

cMyc$^{GOF\text{-}Tg}$ *vs*. WT scRNA Sequencing

cMyc$^{GOF\text{-}Tg}$ or littermate WT mice (*dLckCre*$^{+/-}$) were infected with 4 x $10^6$ PFU LCMV CL13 intravenously. $CD44^+$ $CD8^+$ T cells were sorted from splenocytes on D7 and D15 post-infection on a Sony SH800 sorted. D15 samples were prepared following manufacturer instructions for the Chromium Next GEM Single Cell 3’ Reagent Kit v3.1 (Dual Index). D7 samples were labeled with 10x Genomics CellPlex cell multiplexing oligos, pooled, and prepared following manufacturer instructions for the Chromium Next GEM Single Cell 3’ Reagent Kit v3.1 (Dual Index) with Feature Barcode Technology for Cell Multiplexing. Libraries were sequenced at the UNC High Throughput Sequencing Facility using a NextSeq 2000 P3 flow cell (R1: 28, i1: 10, i2: 10, R2: 90). Gene signature enrichment comparison between Ctrl vs TEX-MYC$^{Unique}$ or Ctrl vs cMyc GOF in Fig. 2N was performed using the R package escape (2.6.1, (*94*)). Gene sets used in this study is available in **Data S3**.

Bulk ATAC-seq

cMyc$^{GOF\text{-}Tg}$ or littermate WT mice (*dLckCre*$^{+/-}$) were infected with 4x$10^6$ PFU LCMV CL13 intravenously. $CD44^+$ PD-$1^+$ $CD8^+$ T cells were sorted from mice 15 days post-infection on a BD FACS Aria II. Samples were prepared following manufacturer instructions using the Active Motif ATAC-Seq kit (Cat# 53150), with each sample receiving a unique combination of i7 and i5 barcodes. Samples were pooled for sequencing on a NovaSeq X Plus 10B flow cell at Admera Health. ATAC processing was done in command line using Bioconda packages: macs2 (v 2.2.9.1), fastqc (v 0.12.1), bwa (v 0.7.18), picard (v 3.1.1), samtools (v 1.19), bedtools (v 2.31.1), and trimmomatic (v 0.39). Adapter sequences were trimmed from reads, using trimmomatic, before mapping to the mm10 reference genome and generating bed files for peak calling, using picard, samtools, and bedtools.

Taiji multiomics-based TF analysis

*Identification of differentially active, less variable, and low-expression TFs.* To analyze TF activity and expression in fig. S1, we performed an integrative analysis of single-cell multiomics data (*29*) using Taiji v2.0, as we recently described (*60*). Counts per million (CPM) of pseudobulk raw gene expression counts were used to calculate the TF activity score (PageRank). Next, we compared the median PageRank across 16 cell states and calculated coefficients of variation (CV) to assess the specificity of each TF in each cell state.

*cMyc regulatory networks in cMyc GOF vs. WT construction.* Differential TF activity in cMyc GOF vs. WT CD8+ T cells was analyzed with Taiji v2.0, integrating scRNAseq and bulk ATAC-seq data from cMycGOF-Tg vs. WT, as described previously (ref: ). TF activity and log2FC were calculated in each context. Subsequently, a cMyc-TF correlation matrix was generated by calculating Spearman’s correlation of edge weights for each TF pair across their common regulatees. From this matrix, we constructed a graphical model using the R package “huge” (ref), which employs the Graphical Lasso algorithm and a shrunken ECDF (empirical cumulative distribution function) estimator. An edge between two TFs was established if their correlation was deemed significant by the model, with a lasso penalty parameter (lambda) of 0.052.

Statistical analyses

For comparisons between two groups, two-tailed unpaired or paired t-tests were performed, with Welch's correction as necessary for significantly different variances. When comparing multiple groups, one-way ANOVA with Tukey's multiple comparison's test was performed. P values $< 0.05$ were considered significant. Unless otherwise noted, error bars are standard error of mean and significance is noted as follows: *, $p \leq 0.05$; **, $p \leq 0.01$; ***, $p \leq 0.001$.

### Figs. S1.

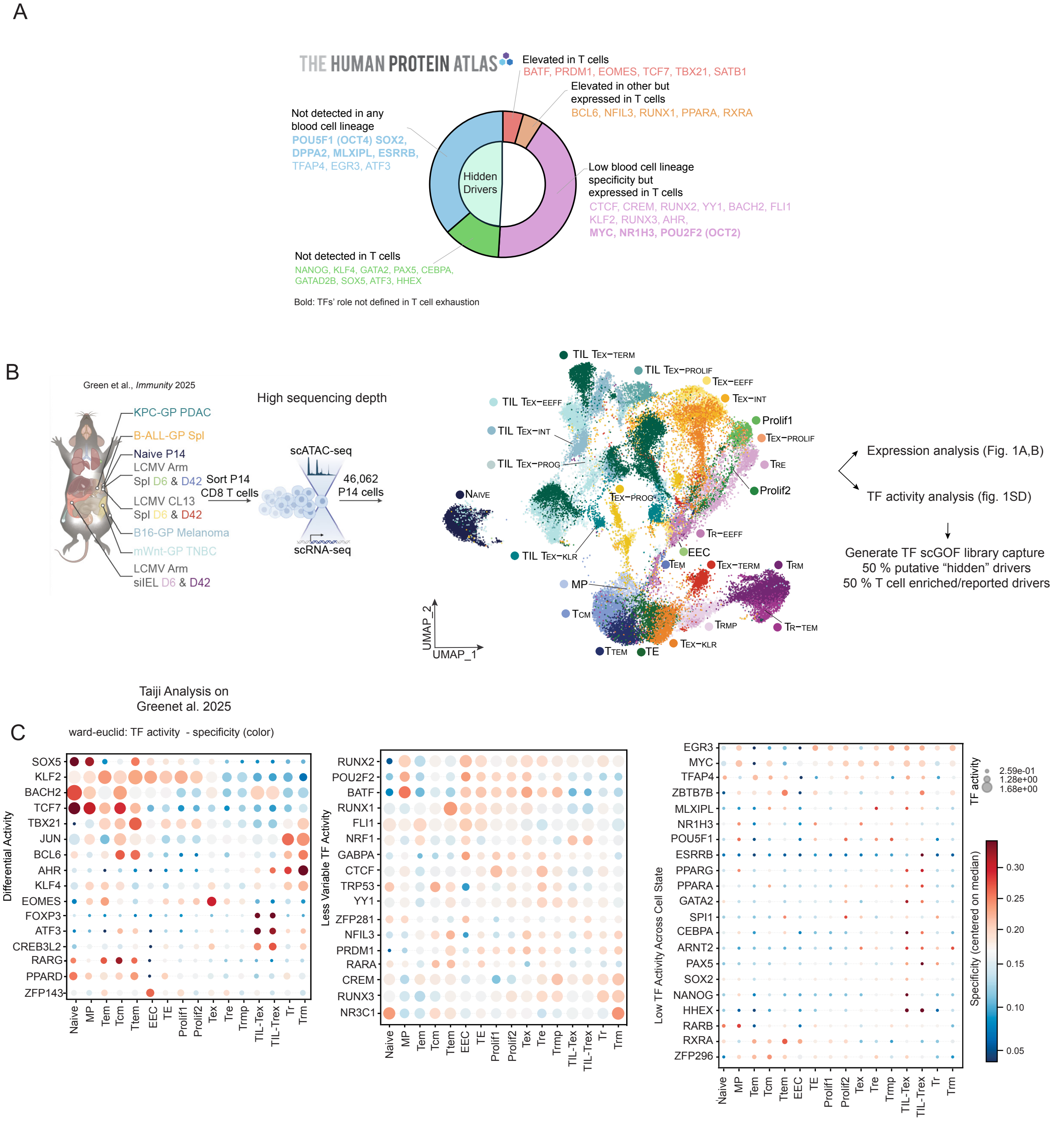


**fig. S1. scGOF-seq framework enables discovery of hidden and underexplored transcriptional regulators.**

**(A)** Classification of TFs based on expression abundance in human T cells using The Human Protein Atlas (*28*). TFs are grouped as enriched in T cells, expressed but not enriched, low blood-lineage specificity but detectable in T cells, or undetectable in T cells. TFs with no defined role in T cell exhaustion are highlighted. **(B)** Integrated single-cell transcriptomic and chromatin accessibility datasets from infection and multiple tumor models used to define T cell state space and TF activity landscapes. **(D)** Bubble plots showing predicted TF activity (color) and expression (size) across CD8$^{+}$ T cell states. Representative TFs are separated by differential, low-variance, or uniformly low activity. **(E)** Schematic of representative ORF retroviral vector used for scGOF-seq library construction.

### Fig. S2.

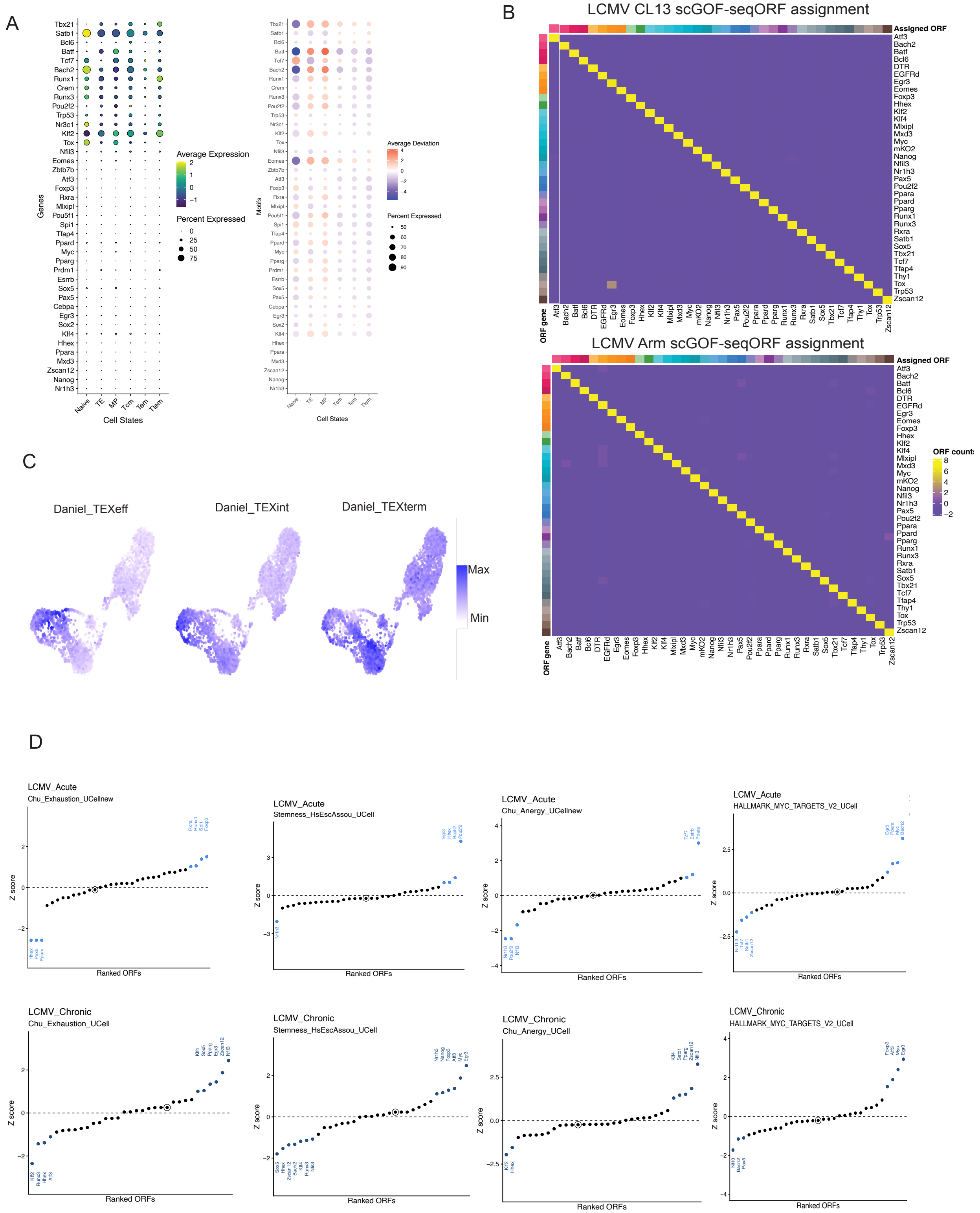


**fig. S2. scGOF-seq performance and TF activity landscapes in CD8⁺ T cells in LCMV infection.**

**(A)** Gene expression analysis in CD8 T cell states in LCMV Arm infection of select TFs included in the scGOF library (left). Chromatin accessibility deviation (ChromVAR) and percent expression of their target genes (right). **(B)** ORF assignment matrix in scGOF-seq demonstrating expression values of each ORF in assigned cells in LCMV Arm and LCMV CL13. **(C)** Gene module enrichment for effector exhausted cells (TEX-EFF), intermediate exhausted cells (TEX-INT), and terminal exhausted cells (TEX-TERM) (*46*) in scGOF-seq screens from **Fig. 1D**. **(D)** Ranked enrichment of ORFs across exhaustion, anergy, stemness, and cMyc-target gene signatures in LCMV Arm and LCMV CL13 scGOF-seq from **Fig. 1D**.

### Fig. S3.

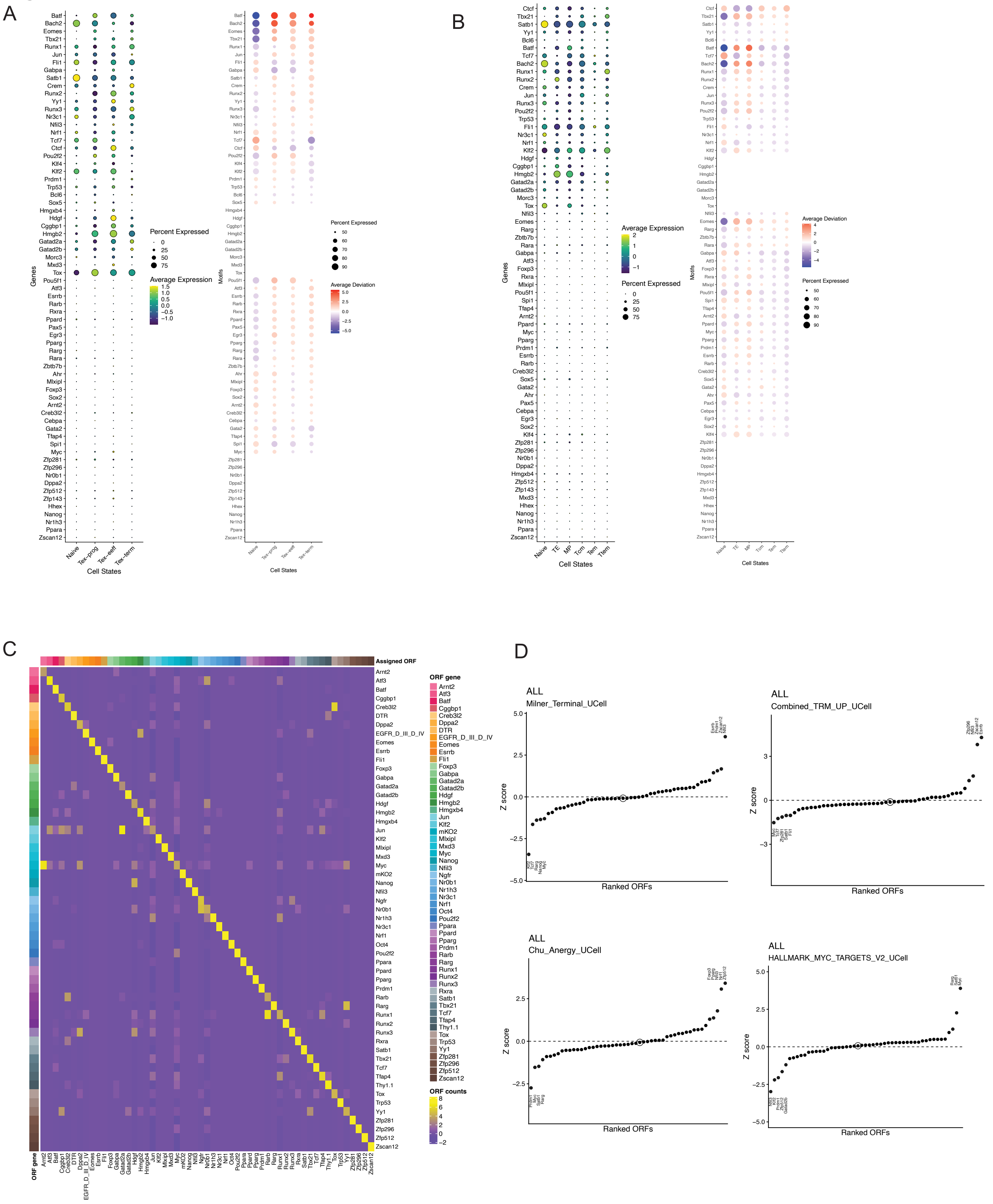


**fig. S3. scGOF-seq identifies TF regulators of CD8$^+$ T-cell differentiation in leukemia.** **(A,B)** Expression and motif accessibility of scGOF library TFs for B-ALL-GP model across CD8$^+$ T-cell states in chronic LCMV CL13 infection (A) and acute LCMV Armstrong infection (B). **(C)** ORF assignment heatmap demonstrating single-cell resolution of pooled GOF perturbations. **(D)** Ranked ORF enrichment across terminal differentiation, anergy, $T_{RM}$, and MYC-target gene signatures.

**Fig. S4.**

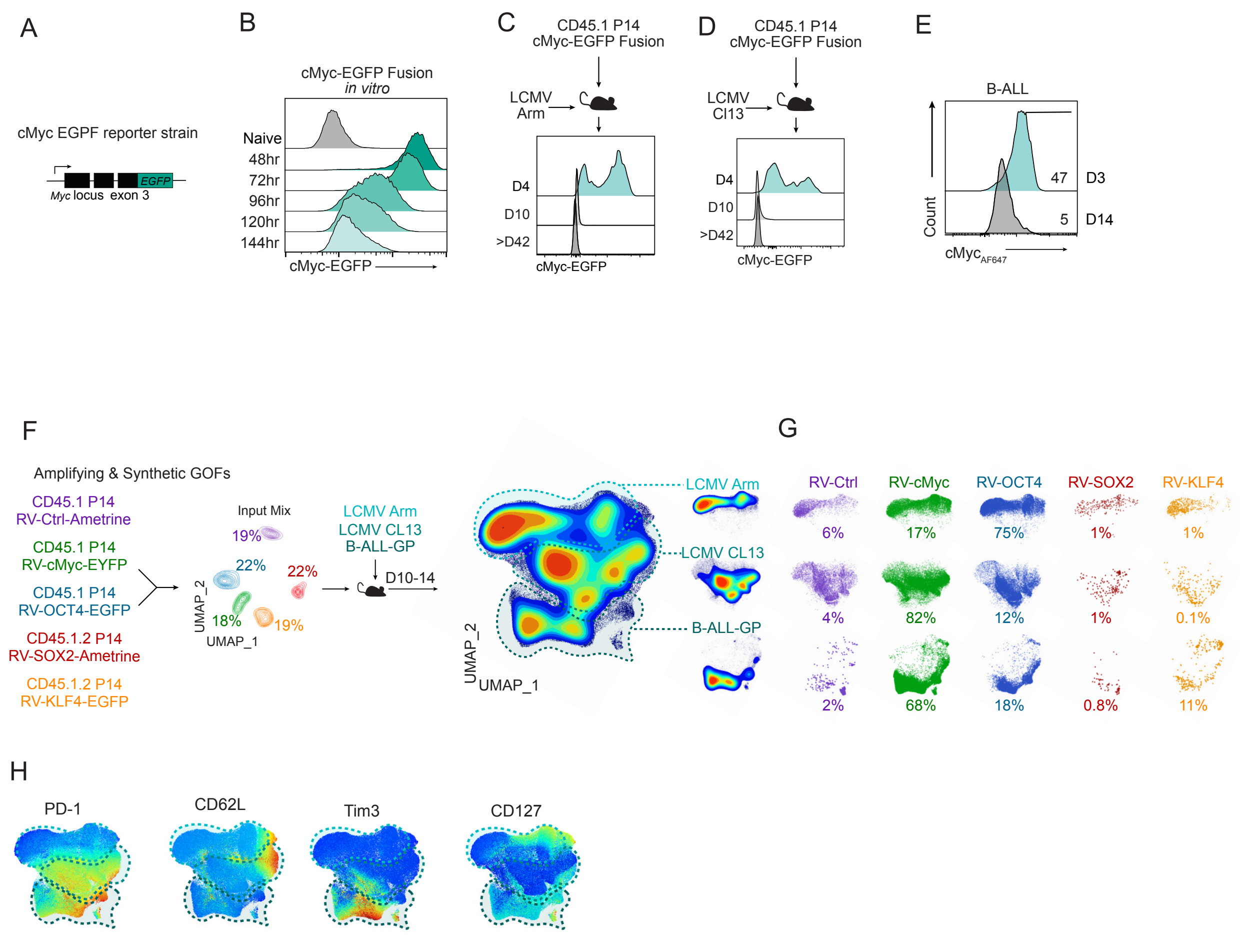


**fig. S4. Hidden driver cMyc and canonical stemness TFs reveals distinctive synthetic GOF phenotypes across immune contexts.**

**(A)** P14 cells from cMyc-EGFP fusion transgenic mice were used to investigate dynamics *in vitro* stimulation and LCMV infection (B-D). **(B-D)** Time-course of cMyc-EGFP expression were tested in T cells isolated from cMyc-EGFP fusion reporter transgenic mice strain following activation *in vitro* (B), acute LCMV Arm infection (C), and chronic LCMV CL13 infection (D). **(E)** cMyc expression dynamics in the B-ALL model analyzed by cMyc antibody staining. **(F)** Experimental design and spectral flow cytometry UMAP projection of five-way mixed transfers of RV-Ctrl, RV-cMyc, RV-OCT4, RV-SOX2, and RV-KLF4 P14 cells in LCMV Arm, LCMV CL13, and B-ALL-GP models. **(G)** Relative contribution of each GOF perturbation across disease contexts. **(H)** Feature overlays for inhibitory and differentiation markers across GOF perturbations.

**Fig.S5.**

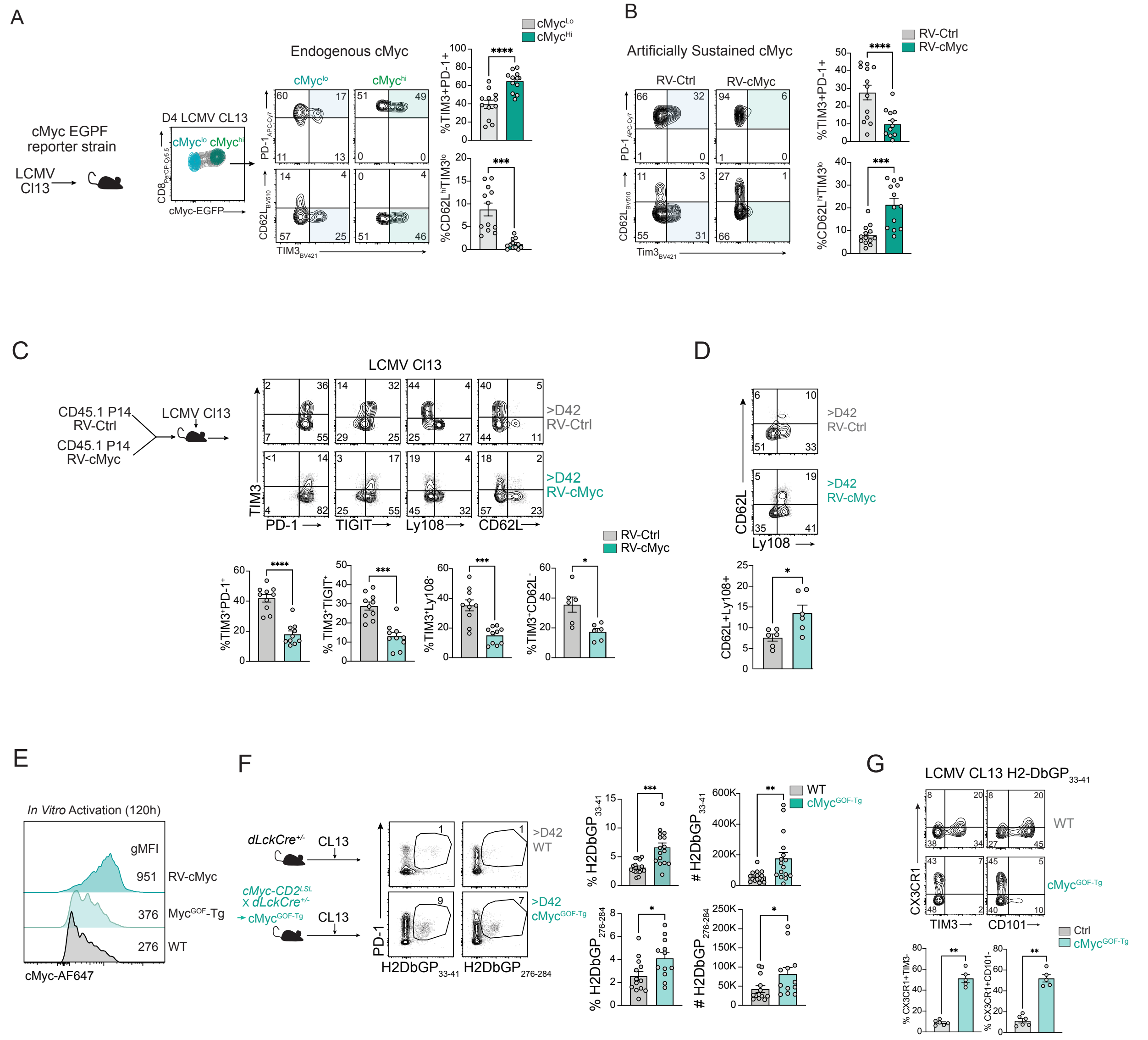


**fig. S5. Unlike endogenous cMyc activity, cMyc GOF limits terminal exhaustion in both TCR-specific and polyclonal CD8$^+$ T cells**

**(A)** cMyc-EGFP fusion reporter mice were infected with chronic LCMV CL13. During early infection (day 4), elevated endogenous cMyc expression is associated with increased frequencies of PD-1$^+$TIM3$^+$ cells and reduced CD62L$^{high}$TIM3$^{low}$ cells. (B) Enforced cMyc expression via RV-cMyc transduction is associated with reduced PD-1$^+$TIM3$^+$ cells and increased CD62L$^{high}$TIM3$^{low}$ cells, in contrast to patterns observed with endogenous cMyc expression in (A). **(C)** Phenotype of RV-Ctrl and RV-cMyc transduced P14 cells at >42 days post LCMV CL13 infection, with quantification derived from the experimental setup shown in **Fig. 2A**. **(D)** Frequency of CD62L$^+$Ly108$^+$ cells from the experiment shown in **Fig. 2A**. **(E)** cMyc expression kinetics following *in vitro* activation of RV-cMyc transduced CD8$^+$ T cells and CD8$^+$ T cells from Myc$^{GOF-Tg}$ mice and wild-type mice. **(F)** WT and Myc$^{GOF-Tg}$ mice were infected with LCMV CL13, and tetramer$^+$ CD8$^+$ T cells were analyzed by spectral flow cytometry at >42 days post-infection. Representative flow plots of H2-DbGP$_{33-41}$ and H2-DbGP$_{276-284}$ tetramer$^+$ cells, with corresponding frequencies and absolute numbers, are shown. **(G)** Representative flow plots and frequencies of CX3CR1$^+$TIM3$^-$ and Ly108$^+$TIM3$^-$ H2-DbGP$_{33-41}$ tetramer$^+$ CD8$^+$ cells. Graphs show mean ± SEM from n = 5–16 mice from one representative experiment or pooled from two or more independent experiments. *$P < 0.05$, **$P < 0.005$, ***$P < 0.001$, paired Student's t-test.

**Fig. S6.**

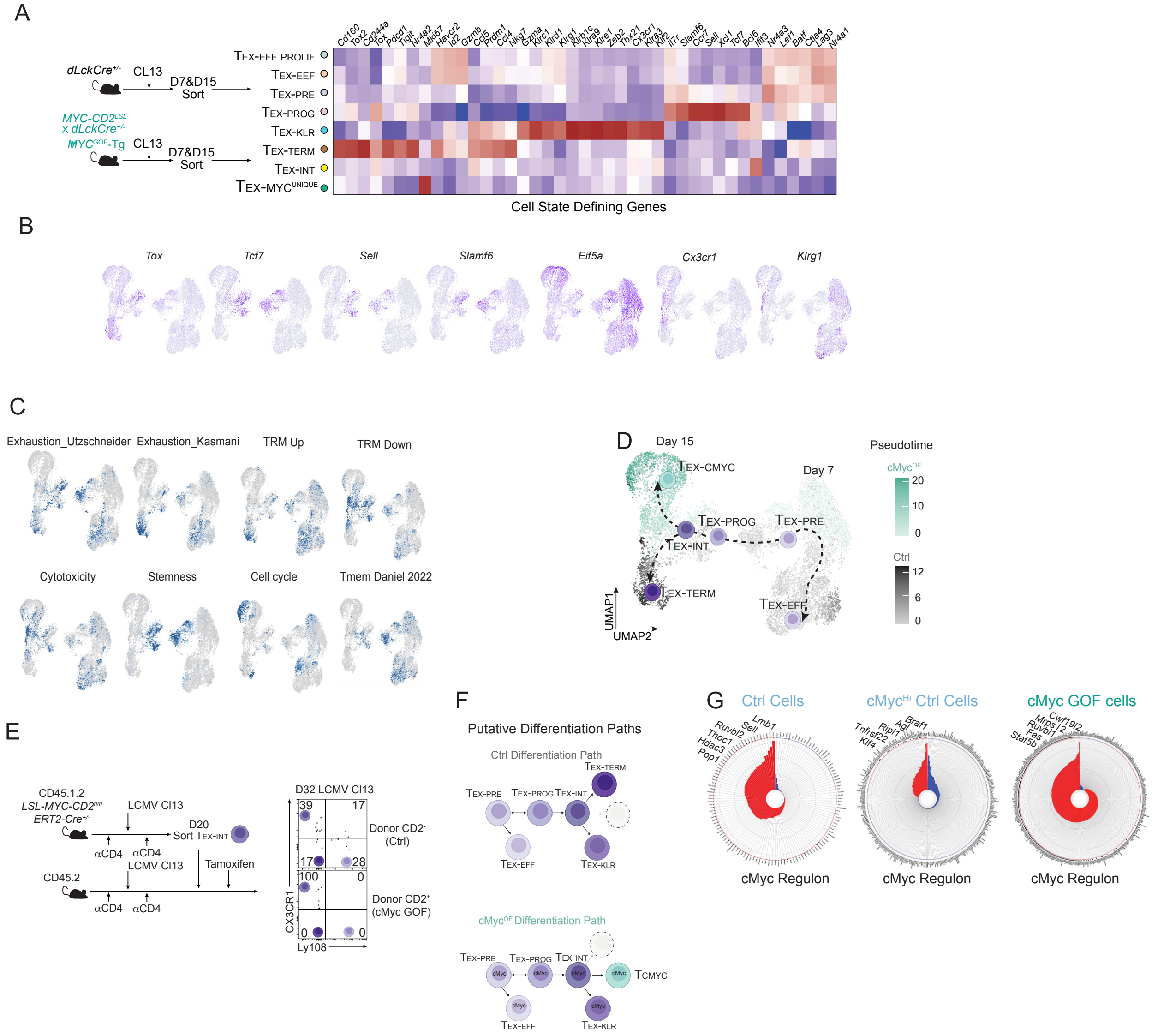


**fig. S6. cMyc GOF redirects exhausted-lineage differentiation into a synthetic trajectory**

**(A)** Expression of representative cell state-defining genes from scRNA-seq analysis of CD8$^+$ T cells isolated from LCMV CL13 infected WT and cMyc$^{GOF\text{-}Tg}$ mice. **(B)** Feature plots of CD8$^+$ T cell differentiation states defining marker gene expression. **(C)** Feature plots of gene-signature scores. **(D)** UMAP projections illustrating the cMyc-GOF induced emergence of unique populations during chronic infection and divergence of differentiation trajectories between WT and cMyc GOF CD8$^+$ T cells. **(E)** Experimental schematic for inducible cMyc GOF activation in sorted TEX-INT cells (PD-1+CX3CR1+Ly108-KLRG1-). TEX-INT cells were from tamoxifen-inducible cMyc$^{GOF\text{-}Tg}$ (CD45.1/CD45.2 MYC-CD2LSL; Cre-ERT2) mice infected with LCMV CL13. The sorted cells were transferred into infection-matched recipient mice. On day 32 of infection, 10 days following tamoxifen treatment, cMyc GOF and Ctrl cells within the same recipient mouse were analyzed for CX3CR1 and Ly108 expression. **(F)** Inferred differentiation trajectories comparing control and cMyc GOF CD8$^+$ T cells. (**G**) scMiner analysis (*30*) to predict context-dependent target genes of cMyc.

**Fig. S7.**

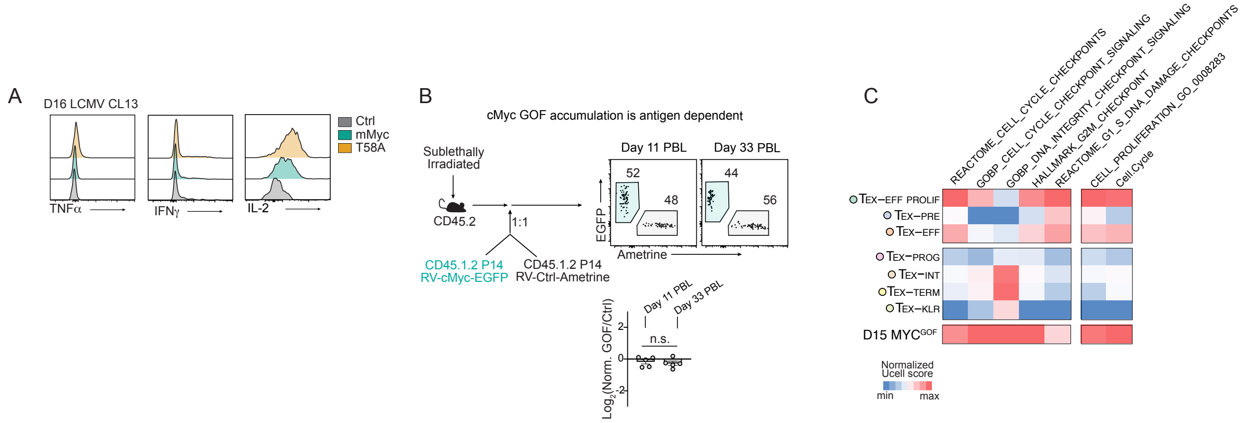


**fig. S7. cMyc GOF promotes antigen-dependent proliferation without loss of cell-cycle control.**

**(A)** Cytokine production (TNFα, IFNγ, IL-2) by Ctrl, cMyc-GOF, and cMycT58A-GOF cells. **(B)** In the absence of antigen, cMyc-GOF–perturbed P14 cells did not show preferential accumulation *in vivo* after mixed transfer, demonstrating that cMyc-GOF–driven accumulation is antigen dependent. **(C)** Gene set enrichment scores for cell-cycle checkpoint pathways across exhausted subsets in day 15 WT and cMyc-GOF cells from LCMV CL13 infection indicate intact checkpoint activity in cMyc GOF cells, with no evidence of compromised cell-cycle regulation.

**Fig. S8.**

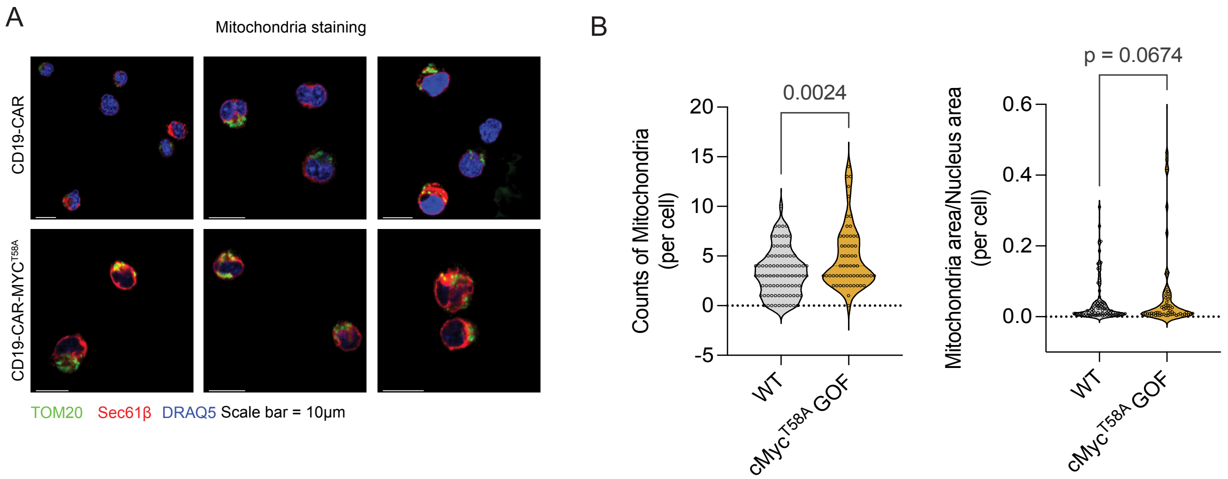


**fig. S8. cMyc GOF increased mitochondira mass in human CAR-T cells. (A)** Confocal microscopy of mitochondrial structure in human anti-CD19 CAR-T cells -/+ MYCT58A GOF cultured in CD19+ Daudi cells. Scale bars indicate 10 µm. TOM20: mitochondria marker, SEC61β: ER marker, DRAQ5: nucleus marker. **(B)** Quantification of mitochondrial counts and area. Data are represented as mean ± SEM. Statistical analysis was performed using unpaired two-sided Student's t test. **p < 0.01, ***p < 0.001, ****p < 0.0001.

### Fig. S9.

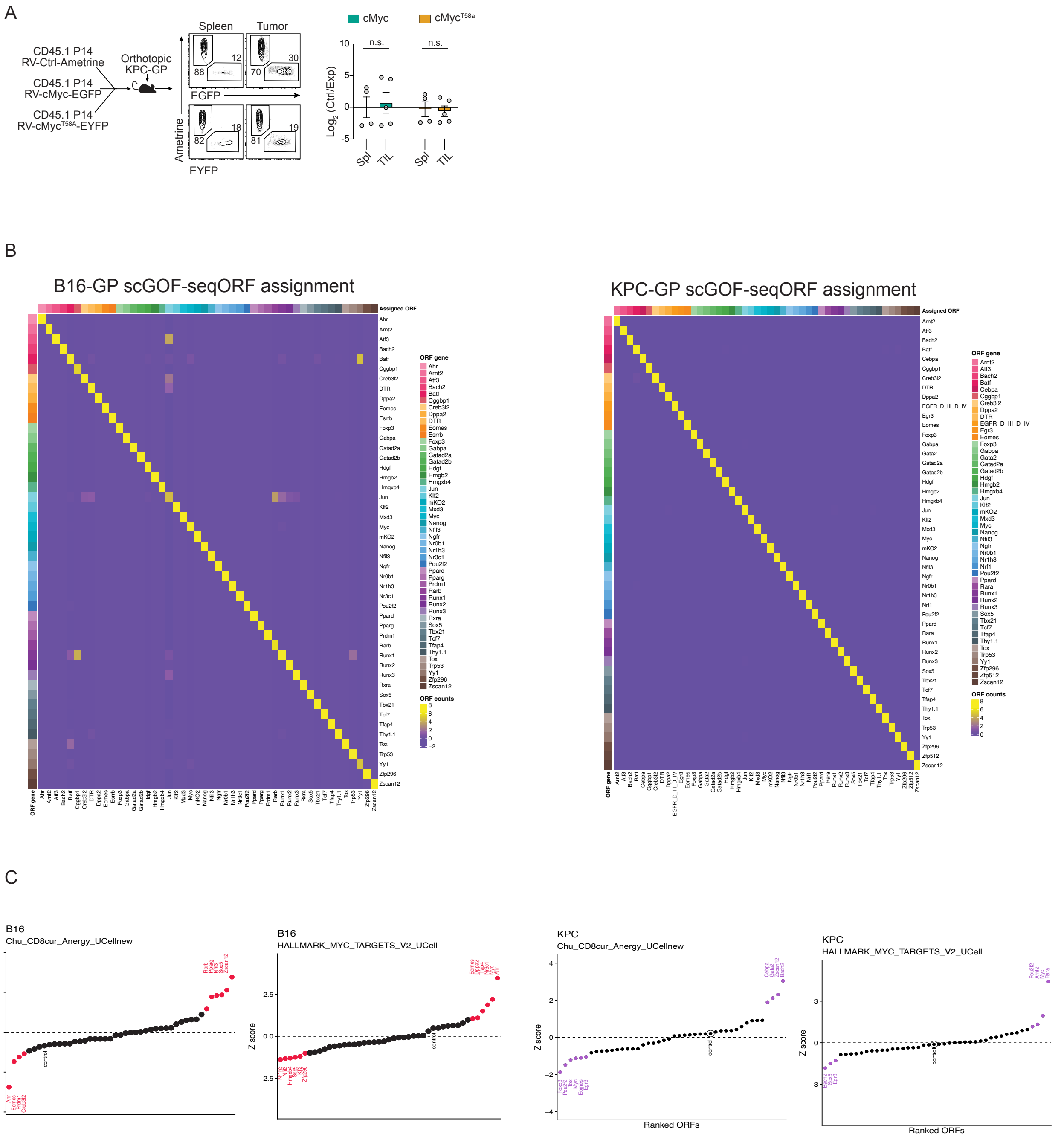


**fig. S9. Second-round scGOF-seq identifies TFs that complement cMyc GOF in solid tumors.**

**(A)** Mixed transfer strategy and relative accumulation of cMyc and cMyc$^{T58A}$ GOF cells in spleen and tumor in KPC-GP models. **(B)** ORF assignment heatmaps for B16-GP and KPC-GP scGOF-seq screens. **(C)** Ranked ORF enrichment across anergy and MYC-target gene signatures.

**Fig. S10.**

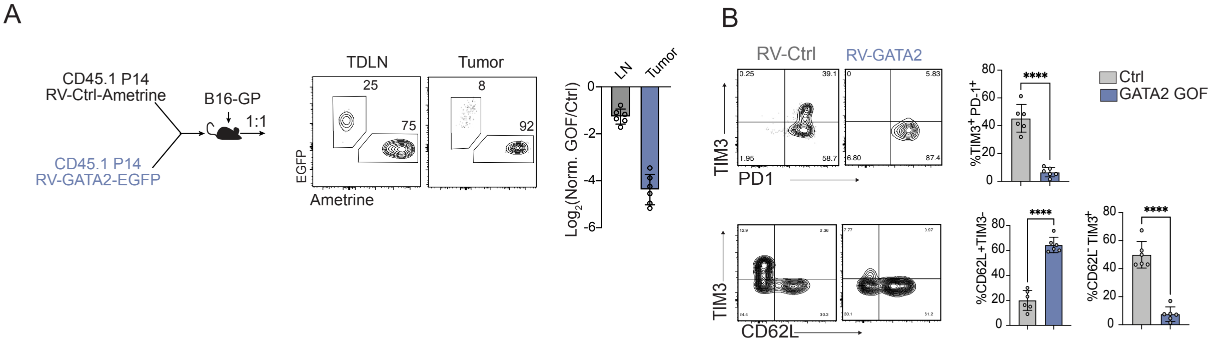


**fig. S10. GATA2 GOF reveals an additional regulator of T-cell differentiation in tumors.** **(A)** Mixed transfer of RV-Ctrl and RV-GATA2 P14 cells into B16-GP tumors and relative accumulation in tumor-draining lymph nodes and tumors. (**B)** Flow cytometric analysis and quantification of PD-1, TIM3, CD62L, CD69, and Ly108 expression following GATA2 GOF.
